# Corticospinal propagation of full-length TDP-43 toxicity drives brain-to-muscle pathology

**DOI:** 10.64898/2026.03.27.714692

**Authors:** Jacopo Marongiu, Valeria Crippa, Isabella Marzi, Clara Porcedda, Maria Cristina Gagliani, Annalisa Brivio, Maria Francesca Palmas, Michela Etzi, Marcello Serra, Maria Antonietta Casu, Ignazia Mocci, Augusta Pisanu, Nicola Simola, Valeria Sogos, Raffaella Isola, Katia Cortese, Alfonso De Simone, Fabrizio Chiti, Anna Rosa Carta

## Abstract

Amyotrophic lateral sclerosis (ALS) is a fatal neurodegenerative disorder characterized by progressive degeneration of upper and lower motor neurons. Cytoplasmic inclusions containing TAR DNA-binding protein 43 (TDP-43), a key regulator of RNA metabolism, represent a pathological hallmark of all sporadic (sALS) and most familial (fALS) forms, underscoring its central role in disease pathophysiology. In affected neurons, full-length (FL) TDP-43 undergoes nuclear-to-cytoplasmic mislocalization, leading to aggregation and cellular dysfunction, and can be released to propagate pathology across neural and non-neural circuits. However, the *in vivo* toxicity and spreading capacity of FL TDP-43 remain poorly defined.

Here, we show that purified, stable human FL TDP-43 was readily internalized by neuronal cells, where it induced aggregation and significantly reduced cell viability. In vivo, an acute unilateral stereotaxic infusion of FL TDP-43 into the rat primary motor cortex was sufficient to trigger a robust centrifugal propagation of pathology along the corticospinal axis and beyond the central nervous system (CNS). TDP-43 pathology spread from the motor cortex to the spinal cord and reached skeletal muscle. At the cellular level, propagated pathology was characterized by intraneuronal phosphorylated TDP-43 (pTDP-43) inclusions, accumulation of high-molecular-weight TDP-43 species, region-specific neurodegeneration, and pronounced mitochondrial vulnerability. Notably, skeletal muscle displayed impaired mitochondrial bioenergetics, accompanied by both motor and non-motor behavioral deficits.

Collectively, our findings demonstrate neuron-to-neuron, brain-to-spinal cord and brain-to-muscle spreading of FL TDP-43 toxicity *in vivo*, establishing a mechanistic link between central TDP-43 pathology and peripheral dysfunction. This work identifies FL TDP-43 as an active driver of disease spreading in ALS and provides the basis for a non-transgenic, TDP-43-driven rat model of disease propagation.

## Introduction

Amyotrophic lateral sclerosis (ALS) is a neurodegenerative disorder characterized by the progressive degeneration of both upper and lower motor neurons, leading to skeletal muscle atrophy, progressive paralysis, and ultimately death, typically within 3 to 5 years from symptom onset.^1,2^ The mechanisms underlying disease initiation and progression remain debated. According to the “dying forward” hypothesis, ALS originates in the motor cortex and propagates along corticofugal projections via anterograde trans-synaptic degeneration.^3^ Conversely, the “dying back” hypothesis posits that pathology begins at the neuromuscular junction or in skeletal muscle and progresses retrogradely toward motor neuron cell bodies.^4,5^ These mechanisms are not mutually exclusive while unifying causal hypothesis has yet to be established, and ALS is increasingly viewed as a multi-system disorder arising from a complex interplay between genetic susceptibility and environmental factors.^6^ Although up to 90% of ALS cases are sporadic (sALS), the familial forms (fALS) have provided critical insight into disease mechanisms.^6^ Among more than 40 genes identified as potential contributors to disease pathogenesis, TAR DNA-binding protein 43 (*TARDBP*), Superoxide dismutase 1 (*SOD1*), Chromosome 9 open reading frame 72 (*C9orf72*), and Fused in sarcoma (*FUS*), are strongly associated with ALS.^7,8^ Notably, the wild-type (WT) protein products of some genes associated with fALS also exhibit aberrant behavior in sALS, including TDP-43 encoded by *TARDBP*, SQSTM1/p62, optineurins, etc. Indeed, cytoplasmic inclusions containing TDP-43 are widely recognized as a pathological hallmark in both sALS and fALS, supporting the pivotal role of this protein in ALS pathophysiology.^9–11^ Moreover, TDP-43 inclusions have been observed across a spectrum of neurodegenerative disorders, including frontotemporal dementia (FTD), limbic-predominance age-related TDP-43 encephalopathy (LATE) and in Perry disease,^12,13^ as well as in Alzheimer’s disease (AD) and Parkinson’s disease (PD).^14–16^

TDP-43 is a ubiquitously expressed RNA/DNA binding protein involved in multiple aspects of RNA-metabolism, including transcription, maturation, transport, and stability.^17^ Although TDP-43 is predominantly localized in the nucleus, it continuously shuttles between the nucleus and the cytoplasm.^17^ In disease states, alterations in specific structural domains, as well as aberrant post-translational modifications such as phosphorylation, ubiquitination, sumoylation, acetylation, citrullination, methylation, and C-terminal cleavage can disrupt the correct nuclear-cytoplasmic distribution.^18–20^ These alterations promote the formation of insoluble cytoplasmic inclusions, ultimately resulting in a loss-of-function and a gain-of-toxic function.^10,17,21–23^ Specifically, phosphorylation and C-terminal cleavage of TDP-43 have been linked to its increased cytoplasmic mislocalization and aggregation.^10,24^ Although full-length (FL) TDP-43 exhibits limited intrinsic aggregation propensity *in vitro*, post-mortem studies have demonstrated that it can form amyloid fibrillar aggregates *in vivo*, contributing to neuronal toxicity.^20,25–27^ Moreover, mitochondrial impairment has been reported in both *in vivo* and *in vitro* models expressing either WT or mutant TDP-43, emerging as a key mediator of TDP-43-induced intracellular toxicity.^28–30^

In addition to intracellular toxicity, increasing evidence supports a prion-like propagation of TDP-43 pathology. Cells counteract toxic protein aggregation through the protein quality-control system (PQC) as well as via extracellular secretion of TDP-43.^31–33^ While protective for the releasing cell, this secretion may promote prion-like spreading: both FL TDP-43 and its C-terminal fragments can be transferred to neighbouring cells, where they seed aggregation of endogenous TDP-43, thereby amplifying pathology across anatomically connected regions.^27,32^ Transgenic animal models harboring either WT or mutant forms of TDP-43 have provided important insights into this process, reproducing the pathological spreading across the central nervous system (CNS).^30,34,35^ However, these approaches rely on overexpression systems, intracortical inoculation of pathological TDP-43 seeds or preformed fibrils,^36,37^ which may not fully recapitulate the behavior of the native protein. In contrast, whether native FL TDP-43 is sufficient to induce toxicity and drive pathological spreading *in vivo* remains unknown. This represents a major gap in the field, largely due to technical limitations in producing stable and functional FL TDP-43, that has posed major challenges for *in vivo* experimentation.^38–41^

Here, we addressed this question by leveraging a robust purification strategy to obtain stable human FL TDP-43 and directly assess its pathogenic potential.^41^ We show that FL TDP-43 was efficiently internalized by neuronal cells, where it induced aggregation, impaired mitochondrial function, and reduced cell viability. Importantly, we demonstrate that the acute unilateral infusion of native FL TDP-43 into the rat primary motor cortex was sufficient to trigger an anatomically defined propagation of pathology along the corticospinal axis. Pathology extended from the motor cortex to the spinal cord and further to skeletal muscle (i.e., gastrocnemius), where it was associated with marked mitochondrial dysfunction and behavioral deficits. At the cellular level, propagated pathology was characterized by intraneuronal phosphorylated TDP-43 (pTDP-43) inclusions, accumulation of high-molecular-weight species, mitochondria abnormalities, and region-specific neurodegeneration. These findings provide direct *in vivo* evidence that FL TDP-43 is an active driver of toxicity and disease propagation. Collectively, this study establishes the intrinsic pathogenicity of native FL TDP-43 *in vivo* and identifies a brain-to-muscle axis of disease spreading. Moreover, it introduces a non-transgenic, TDP-43-driven rat model that recapitulates key features of ALS.

## Materials and methods

### FL TDP-43 expression and purification

Human FL TDP-43 cloned into a pNic28-Bsa4 plasmid was recombinantly expressed and purified as previously described.^42^ In brief, *Escherichia coli* BL21 (DE3) cells transformed with the plasmid were grown overnight at 37 °C in Luria-Bertani (LB) medium (20 g/L) supplemented with 50 µg/ml kanamycin. The overnight culture was diluted into fresh LB and further incubated at 37 °C until reaching an OD_600_ of 0.8, after which expression was induced overnight at 37 °C with 1 mM isopropyl β-D-1-thiogalatoside (IPTG). Bacteria cells were harvested by centrifugation at 6,840 *g*, for 30 min, at 4 °C and the pellet was resuspended in lysis buffer (50 mM Tris-HCl, 0.5 M NaCl, pH 8.0). The resulting suspension was sonicated on ice at 20 kHz for six cycles of 20 s, 100% amplitude, and then centrifuged at 20,700 *g*, for 30 min, at 4 °C to separate the pellet containing the inclusion bodies (IBs) from the supernatant. Subsequently, the IBs were washed in three steps, each followed by centrifugation at 20,700 *g*, for 30 min, at 4 °C, using the following buffers: (1) 50 mM Tris-HCl, 1% (v/v) TritonX-100, pH 8.0; (2) 50 mM Tris-HCl, 2 M NaCl, pH 8.0; (3) 50 mM Tris-HCl, pH 8.0. After the third centrifugation, the washed IBs were solubilized overnight at room temperature under stirring in buffer A (50 mM Tris-HCl, 8 M urea, 0.5 M NaCl, 10 mM imidazole, 1 mM DTT, pH 8.0). The solubilized IBs were then centrifuged at 20,700 *g* for 30 min, at 4 °C, and the supernatant containing TDP-43 was loaded onto a 10 ml of pre-equilibrated HisPur Ni-nitrilotriacetic acid (Ni-NTA) resin (Thermo Fisher Scientific) in a gravity flow column. The Ni-NTA resin was washed with buffer B (50 mM Tris-HCl, 8 M urea, 0.5 M NaCl, 25 mM imidazole, pH 8.0) and TDP-43 was eluted using buffer C (50 mM Tris-HCl, 8 M urea, 0.5 M NaCl, 250 mM imidazole, pH 8.0). Refolding of TDP-43 was carried out by a 20-fold slow dropwise dilution at 4 °C into a refolding buffer containing 50 mM HEPES, pH 8.0, 9.6 mM NaCl, 10 mM KCl, 2 mM CaCl_2_, 2 mM MgCl_2_, 0.8 M sucrose, 0.5 M L-arginine, 0.6 M guanidine, 0.5% (v/v) Triton X-100, 0.2% (w/v) PEG 3350, 1 mM reduced glutathione (GSH), 0.1 mM glutathione disulphide (GSSG), and one tablet of EDTA-free Pierce Protease Inhibitor (Thermo Fisher Scientific). The resulting solution was concentrated at 4 °C by ultrafiltration using a 400 mL Amicon stirred cell (Millipore Sigma) with a 10 kDa molecular weight cut-off (MWCO) membrane filter. The concentrated sample was then loaded at 4 °C onto a HiLoad 16/600 Superdex 200 pg column pre-equilibrated with SEC buffer (50 mM HEPES, 0.2 M KCl, 0.1 M imidazole, 0.1 % (w/v) PEG 3350, 0.1 M sucrose, 0.025 % (v/v) TritonX-100, pH 8.0) and eluted isocratically at 1 mL/min using an AKTA Pure 25 L system (GE Healthcare, Wakesha, WI). Finally, the TDP-43 containing fraction was subjected to an ion exchange chromatography using the same AKTA Pure 25 L system and a HiTrap Q Sepharose Fast Flow (QFF) 5 mL column pre-equilibrated with buffer A QFF (50 mM HEPES, 0.1% (w/v) PEG 3350, 2 mM dodecyldimethylamine oxide (LDAO), pH 8.0) and eluted using a step-gradient in buffer B QFF (50 mM HEPES, 0.1% (w/v) PEG 3350, 2 mM (LDAO), 0.25% (w/v) octyl-β-D-glucopyranoside (OG), 1 M NaCl, pH 8.0). Purified TDP-43 contained 414 residues with the addition of the MHHHHHHSSGVDLGTENLYFQS sequence fused to the N-terminus before Met1, for a total of 436 residues. It was stored at concentrations ranging from 0.9 to 1.4 mg/mL (20 – 30 µM) in buffer B QFF at −20 °C. Protein concentration was determined with a Shimadzu UV-1900 UV−vis spectrophotometer, with an extinction coefficient at 280 nm (ε_280_) of 46,785 and a molecular weight of 47,292.53 Da. Protein quality control was assessed with SDS-PAGE for purity, intrinsic fluorescence for packing, far-UV circular dichroism for folding, and dynamic light scattering for size, as previously reported.^41,42^ The Limulus Amebocyte Lysate (LAL) test was carried out to exclude the presence of any liposaccharide endotoxin in the TDP-43 solution before biological assays were applied.

### Cell culture and treatments

The human SH-SY5Y neuroblastoma cell line was grown in high glucose DMEM supplemented with 10% fetal bovine serum, 100 units/mL penicillin and 100 µg/mL streptomycin and kept at 37 °C in a humid environment containing 5% CO2 and 95% air.

### TDP-43 internalization

Human FL TDP-43 was thawed and centrifuged at 18,000 *g*, 15 min, at 4 °C, and the supernatant was labelled with the green, fluorescent CF488A dye (# SCJ4600016, Sigma Aldrich, St.Louis, MO, USA). The dye was incubated with TDP-43 in a molar ratio dye:protein of 1:2 and the reaction was carried out by stirring the mixture overnight at 4 °C; thereafter the unbound dye was removed using ultra-centrifugal filter with 10 kDa cutoff (#UFC201024 Merck KGaA, Darmstadt, Germany) and the concentration was measured using a nanophotometer. Protein internalization was evaluated in SH-SY5Y cells after 24 hours exposure to 0.12 µM CF488A-labelled TDP-43. Nuclei were counterstained with Hoechst 33258. Fluorescence images were acquired using Olympus BX41 fluorescence microscope at a 60X magnification.

### Cell viability

Cell viability was assessed by MTT assay. SH-SY5Y cells were plated at a density of 1× 10^4^ cells/well in 96-well plates and incubated for 2, 6 or 24 hours with 0.12 μM human FL TDP-43. After treatment, MTT reagent (0.5 mg/mL) was added for 3 hours at 37 °C. The resulting formazan crystals were dissolved in DMSO and absorbance was measured at 570 nm using a spectrophotometer (EnVision plate analyzer, Perkin Elmer, Milan, Italy).

### Mitochondrial membrane potential

To assess mitochondrial membrane potential, SH-SY5Y cells were seeded in 96-well plates at a density of 1× 10^4^ cells/well and treated with 0.12 μM human FL TDP-43 for 2, 6 or 24 hours. Afterwards, cells were stained with JC-1 dye (ab113850, Abcam, Cambridge, UK), according to the manufacturer’s instructions. JC-1 accumulates in mitochondria in a membrane potential–dependent manner, shifting fluorescence from red to green in depolarized (apoptotic) mitochondria. Fluorescence intensity was quantified using a spectrophotometer setting excitation/emission wavelengths at 535/590 nm to detect red JC-1 aggregates and 488/535 nm to detect JC-1 green monomers.

### Animals and stereotaxic surgeries

All procedures were performed in accordance with the ARRIVE guidelines and in accordance with the guidelines and protocols approved by the European Community (2010/63UE L 276 20/10/2010). Experimental protocols were approved by the Italian Ministry of Health (authorization N 657/2022-PR). All efforts were made to minimize animal pain and discomfort and to reduce the number of experimental animals used.

In order to find the optimal volume and concentration of FL TDP-43, male Sprague Dawley rats (275–300 g, Envigo) (N= 6), were deeply anesthetized with Fentanyl (3 mg/kg) and medetomidine hydrochloride (0.35 mg/kg) and positioned in a stereotaxic apparatus (DKI-900LS, David Kopf Instruments, Tujunga, CA). Rats were then bilaterally infused with either a fluorescently-labelled (Alexa Fluor 488 C5-Maleimide) FL TDP-43 at three di different concentrations (5 μM; 10 μM; 15 μM) or vehicle (TDP-43 buffer B QFF: 50mM Hepes, 2mM LDAO, 0.1% PEG3350, 1 M NaCl, 0.25% OG, pH 8.0) into the primary motor cortex (coordinates relative to bregma; +1.9 mm anteroposterior; ± 2.8 mm mediolateral; −1.6 mm beneath the dura) according to the atlas of Paxinos and Watson.^43^ Stereotaxic coordinates were chosen according to a previous study.^44^ One day after surgery, rats were deeply anesthetized and transcardially perfused with ice-cold 0.1 M PBS (pH 7.4), followed by 4% buffered paraformaldehyde, the brains were collected and vibratome-cut in 40 μm slices containing the motor cortex. Slices were then observed under a fluorescence microscope (Zeiss, 5X) to verify the infusion site, to detect the extent of FL TDP-43 diffusion within the motor cortex or along the injector trace, as well as the deposition of fluorescent aggregates (data not shown). The final infusion volume of 5 μL with 10 μM concentration was selected.

Thereafter, a subsequent group of rats (N=45) were deeply anesthetized with Fentanyl (3 mg/kg) and were homolaterally infused with 5 μL of FL TDP-43 (N= 22; 10 μM) or vehicle (N= 23) into the primary motor cortex (coordinates relative to bregma; +1.9; 0 mm anteroposterior; ± 2.8; ± 2 mm mediolateral; −1.6 mm beneath the dura) at the infusion rate of 1 μL/min via a silica microinjector. The injector was left in place for an additional 5 min after the infusion was completed, then gradually withdrawn to prevent backflow along the needle track.

### Immunofluorescence

Four months after FL TDP-43 or vehicle infusion, and one day after completion of behavioral testing, half of the rats (N = 23) were anesthetized and transcardially perfused with ice-cold 0.1 M PBS (pH 7.4), followed by 4% buffered paraformaldehyde. This time point was chosen based on previous evidence from our laboratory showing that one month after infusion, TDP-43 produced no effect on either mitochondrial complex IV (CIV) expression levels in cortical neurons or motor performance (Fig. S6). After perfusion, the brain and the spinal cord were carefully removed, post-fixed overnight in 4% paraformaldehyde and stored in 0.1% NaN3-PBS at 4 °C. Brain cortex was vibratome-cut in 40-μm-thick serial coronal sections. Spinal cord was removed from the NaN3-PBS solution and stored in 30% sucrose solution for one week. Thereafter, spinal cord was cryostat-cut in 20 μm-thick serial coronal sections.

Sections were preincubated with a blocking solution with normal serum/BSA and then were immunoreacted with the following unconjugated primary antibodies for double immunolabeling: mouse monoclonal anti-NeuN (1:500, Abcam); rabbit polyclonal anti CIV (1:100, Thermo Fisher Scientific); rabbit monoclonal anti pTDP-43 (1:500, Thermo Fisher Scientific).

For fluorescence visualization of NeuN and CIV a two-step indirect labelling protocol was used (Alexa Fluor 594 anti-mouse 1:500, Jackson ImmunoResearch West Grove, PA, USA; Alexa Fluor 488 anti-rabbit 1:500 Jackson ImmunoResearch, West Grove, PA, USA), whereas a three-step detection was carried out to increase the signal of pTDP-43 by combining biotin-SP-conjugated IgG (1:500, Jackson ImmunoResearch West Grove, PA, USA) and streptavidin–fluorescein (1:400, Jackson ImmunoResearch), as previously described (Boi et al., 2020; Palmas et al., 2022)(Boi et al., 2020; Palmas et al., 2022). Images were acquired using a spinning disk confocal microscope (Crisel Instruments) with a X63 magnification.

### Confocal Microscopy Analysis

Qualitative and quantitative analyses for NeuN, pTDP-43, CIV were performed using a spinning disk confocal microscope (Crisel Instruments, Rome, Italy) with a ×63 magnification. Surface rendering, colocalization, maximum intensity, and simulated fluorescence process algorithms were used (ImageJ and Imaris 7.3). A stack was obtained from each dataset (40 images). In the resulting stacks, for each animal, four regions of interest covering the primary motor cortex on each acquired section were chosen (x = 1024 μm; y = 1024 μm; z = 40 μm), and the volume of the elements was calculated. For colocalization analysis, a colocalization channel was automatically generated by Imaris 7.3.

In the resulting stacks, a subset of randomly chosen NeuN+ cells was identified and selected, and the colocalized volume of CIV was calculated. Mean colocalization values obtained from cells analyzed in each animal from each experimental group were plotted as a frequency distribution displaying the percentage of colocalization between CIV signal and the selected NeuN+ cell. Frequency distribution analysis indicated a different CIV expression across experimental groups. For this reason, an appropriate cut-off value was set to categorize the identified cell populations into high and low expressing CIV cells; mean values within each class were calculated for each experimental group and statistically compared.

### Electron Microscopy and Mitochondria Analysis

Slices from brain cortex and spinal cord were washed in cold PBS1X and then fixed for 3 hours with 2.5% glutaraldehyde in 0.1 M Na+Cacodylate buffer at room temperature. Slices were post-fixed in 4% osmium tetroxide (cat.19140, Electron Microscopy Science, Hatfield, PA, USA) for 2 hours and in 1% aqueous uranyl acetate (cat. 22400-1, Electron Microscopy Science) for 1 hour at room temperature. Slices were then dehydrated through a graded ethanol series with propylene oxide as a transition fluid (cat. P021, TAAB Laboratories Equipment, Aldermaston, UK) and embedded in epoxy resin (Poly-Bed; cat. 08792-1, Polysciences, Warrington, PA, USA) overnight at 42°C e ultimately for 2 days at 60 °C. Subsequent examination of the regions of interest (FL TDP-43 infused regions of cortex and spinal cord) was performed on 200 nm semithin sections stained with Toluidine Blue. Ultrathin sections (50 nm) were then cut and counterstained with 1% uranyl acetate. Electron micrographs were acquired using a HT7800 120 kV transmission electron microscope equipped with Megaview III digital camera and Radius 2.0 software (EMSIS, Muenster, Germany). For ultrastructural morphometry, three independent animals per condition were analyzed. In each animal, 10 morphologically identifiable neurons from the region of interest were selected, and all mitochondrial profiles visible within the soma in the acquired ultrathin sections were included in the analysis. Mitochondrial length was measured on TEM images using the line tool in Radius 2.0 software, and the number of mitochondrial profiles per neuron/field was also recorded. Quantitative values were averaged per animal and used for statistical analysis.

### Protein Extraction from frozen tissues

To extract total proteins, frozen motor cortex, cervical spinal cord, and gastrocnemius muscle were pulverized using a pre-chilled stainless-steel mortar and pestle. 20 mg of pulverized tissues were then homogenized using a Tissue-Lyser II (Qiagen, Hilden, Germany) with stainless steel beads in 200 μl of RIPA lysis buffer (150 mM NaCl, 6 mM Na₂HPO₄, 4 mM NaH₂PO₄, 2 mM EDTA pH 8.0, 1% sodium deoxycholate, 0.5% Triton X-100, 0.1% SDS), supplemented with a protease inhibitor cocktail (Complete tablets, Roche Diagnostics GmbH, Mannheim, Germany, 04693116001) and phosphatase inhibitors (100 mM sodium orthovanadate and 100 mM sodium fluoride). Lysates were clarified by centrifugation at 10,000 × g for 10 minutes at 4 °C to remove insoluble material. Protein concentration in the supernatants was measured using the bicinchoninic acid (BCA) assay (Cyanagen Reagents for Molecular Biology, Bologna, Italy, PRTD1,0500).

### Western Blot Analysis on motor cortex, spinal cord and gastrocnemius homogenates

Western blot analysis of motor cortex, spinal cord, and gastrocnemius samples was conducted by loading 20 μg of total tissue proteins per lane onto 12% SDS-PAGE gels. After separation, proteins were transferred onto 0.45 μm nitrocellulose membranes (Amersham™ Protran™ Premium, Cytiva, 10600003) using the Trans-Blot® Turbo™ transfer system (Bio-Rad Laboratories, Hercules, CA, USA, 1704150). Membranes were blocked for 1 hour in 5% non-fat dry milk (PanReac AppliChem ITW Reagents, Darmstadt, Germany, A0830,0500) diluted in TBS-T buffer [20 mM Tris base, 140 mM NaCl, pH 7.6, and 0.1% Tween 20 (Merck, Darmstadt, Germany, P1379)] and incubated overnight at 4 °C with primary antibodies: anti-GAPDH (Immunological Science, Roma, Italy, MAB-10578, 1:5,000), anti-TDP-43 C-term (Proteintech, Sankt Leon-Rot, Germany, 12892-1-AP, 1:1,000), anti Phospho-TDP43 (Ser409/410) (pTDP-43) (Proteintech, 66318-1-IG, 1:1,000), anti-HSP70/HSC70 (Santa Cruz Biotrechnology, Santa Cruz, CA, USA, sc-24, 1:1,000), anti-SQSTM1/p62 (Merck, P0067, 1:3,000), anti-HSPB8 (Thermo Fisher Scientific, Waltham, MA, USA, PA5-76780, 1:1,000), and anti-BAG3 (Abcam, Cambridge, United Kingdom, ab47124, 1:3,000), anti-LC3 (Merck, L8918, 1:3,000). Immunodetection was performed using the appropriate HRP-conjugated secondary antibodies: goat anti-rabbit (Jackson ImmunoResearch, 111–035-003, 1:10,000) or goat anti-mouse (Jackson ImmunoResearch, West Grove, Pennsylvania, USA, 115–035-003, 1:10,000). Signal was developed using the enhanced chemiluminescence reagent Westar Antares (Cyanagen, XLS0142) or Westar ETA C ULTRA 2.0 (Cyanagen, XLS075), and images were acquired using the ChemiDoc XRS+ system (Bio-Rad Laboratories, 1708265). Band intensities were quantified using Image Lab software version 5.2.1 (Bio-Rad Laboratories).

### Filter Retardation Assa**y** on motor cortex, spinal cord and gastrocnemius homogenates

A total of 10 μg of motor cortex, spinal cord and gastrocnemius proteins was diluted in 100 μl of RIPA buffer and loaded to a dot-blot apparatus (Bio-Rad Laboratories, 1703938) for filtration through a 0.22 μm cellulose acetate membrane (Whatman, Maidstone, United Kingdom, 00404180). The proteins were fixed with 20% methanol, and the membranes were blocked for 1 hour in TBS-T containing 5% non-fat dry milk. Subsequently, membranes were incubated overnight with an anti-TDP-43 C-terminal antibody (Proteintech, 12892-1-AP, 1:1,000 dilution). After washing with TBS-T, membranes were incubated with a peroxidase-conjugated anti-rabbit secondary antibody (1:5,000). Signal detection was performed following standard Western blot procedures.

### Bioenergetics of isolated mitochondria from the gastrocnemius muscle

Mitochondria were isolated from the left and right gastrocnemius muscles using differential centrifugation. The gastrocnemius was selected for this study because it predominantly contains red muscle fibers, which yield suitable quantities of mitochondria for oxygraphic analysis. The right gastrocnemius resulted as contralateral to the infusion site (with higher probability of being influenced by TDP-43), while the left gastrocnemius was ipsilateral and, presumably, unaffected. Mitochondrial isolation followed previously published methods.^47,48^

Mitochondrial bioenergetics was evaluated at 37°C with a Clark-type electrode (Oxytherm, Hansatech Instruments Ltd., Norfolk, United Kingdom). Mitochondria were tested at a concentration of 0.25-0.125 mg/ml, in a respiration buffer (125 mM KCl, 20 mM Tris Base, 1 mM EGTA, pH 7.2 supplemented with 0.2 % BSA and 10 mM Pi).

The following substrates were used for oxidative phosphorylation (OXPHOS): 10 mM malate/20 mM Glutamate (Complex I, CI), 7.5 µM Rotenone/20 mM Succinate (Complex II, CII), palmitoyl carnitine 40 µM/5 mM malate (lipid substrates), 7.5 µM Rotenone/2.5 mM Ascorbate/ 0.5 mM TMPD and 5 mM Ascorbate (Complex IV, CIV). To these substrates 0.2 or 2 mM ADP were added to stimulate OXPHOS and 2 µg/mL oligomycin or 0.2 mM DNP to assess respectively state 3, high ADP, state 4, or uncoupled respiration. All values reported refer to the final concentrations.

### Western Blot of isolated mitochondria from the gastrocnemius muscle

Western blot analysis was performed on isolated mitochondria (both from contralateral and ipsilateral gastrocnemius). Mitochondrial protein concentration was measured using Pierce^TM^ BCA Protein Assay Kit (#23227, ThermoFisher, Waltham, MA, USA). Subsequently, samples were diluted in Pierce^TM^ LDS non-reducing loading buffer (#84788, ThermoFisher, Waltham, MA, USA) and kept at room temperature to avoid protein aggregation induced by heating. Sixty micrograms of proteins were run on 4-12% Bis-Tris gels (#MP41G10, Millipore Merck, Darmstadt, Germany) at 130 V and transferred onto PVDF membranes (Immobilon E #IEVH00005, Millipore Merck, Darmstadt, Germany) for 1 hour at 100 V. Membranes were blocked with 5% non-fat dry milk for 3 hours at room temperature and incubated at 4°C overnight with anti OxPhos antibody Cocktail (1:500, **#** 45-8099, ThermoFisher, Waltham, MA, USA). The day after, membranes were washed with TBS Tween 0.1% and incubated for 1 hour at room temperature with horseradish-peroxidase-conjugated anti-mouse IgG (1:10,000; #115-035-003 Jackson ImmunoResearch, Baltimore Pike, PA, USA). Consequently, bands were developed with a chemiluminescent substrate (Clarity ECL #1705060, Biorad, Hercules, CA, USA) and visualized with Image Quant LAS-4000 (GE Healthcare, Little Chalfont, UK). Notably, each mitochondrial complex exposure time was optimized to enable accurate densitometric analysis. The signal was normalised with total proteins detected with Coomassie brilliant blue R 0.025% in 40% methanol, 7% acetic acid for 5 minutes. Band intensity was quantified using Image Studio software (LI-COR, Biosciences).

### Behavioral Tests

Behavioral studies were conducted over a 4-day interval, beginning 15 weeks after TDP-43 or vehicle infusion. Prior to each test, rats were acclimated to the testing room for 30 min in order to avoid any alteration in behavioral parameters induced by the novel environment. Tests were carried out between 9 am and 3 pm. All tests were performed and analyzed by individuals blinded to the experimental conditions.

### Beam Challenging test

the challenging beam test was used to assess motor coordination and balance.^45,46^ The apparatus consisted of a 2-meter wooden beam inclined at 15°, connecting a starting platform elevated 40 cm above the floor to the home cage. Three beam widths were tested: 15 mm, 10 mm, and 5 mm. Rats were trained for 3 consecutive days prior to testing.

On the test day, each rat was placed at the lower end of the beam and video recorded while crossing toward the home cage. The number of stepping errors was recorded for each beam width.

Whether an animal failed to complete the task within the set time or fell from the beam, a numerical increment to the error score was applied as follows: +0.25 for 75% beam completion, +0.5 for 50%, and +0.75 for 25%.^46^

### Grip strength test

the grip strength test was used to assess muscular strength and fatigue as demonstrated by Meyer et al.^49^ with a protocol adapted from.^50–52^ Muscle strength and fatigue were measured using a strength grip meter (BIO-GS4 Bioseb, Vitrolles, France). Briefly, rats were permitted to grip the grid with limbs. Rats were then pulled gently back until the grip was released. Each rat was subject to two trials of five consecutive measurements with a resting period of one minute between each trial in their home cage (first test). The same procedure was performed three hours later (second test). The values of each trial were average to calculate the grip strength. A decrease in grip force after the second trial or a lack of recovery between the first and second tests were interpreted as muscular fatigue. All the measurements were normalized to the rat body weights.

### Ultrasonic vocalizations

experiments were performed in a quiet room under an illumination of 40 lx. For recordings, two unacquainted rats paired for condition (vehicle, 5 dyads; TDP-43, 5 dyads) were placed in Plexiglas cylinders (test cages) (diameter, 25 cm; height, 30 cm) having one half completely painted black and the other half partially painted black to create alternating horizontal black and transparent stripes, and the bottom covered with sawdust. Each cylinder was topped with a lid equipped with an ultrasonic microphone (CM16/CMPA, Avisoft, Berlin, Germany) that was connected to an ultrasound recording device (Ultrasound Gate 116 Hb, Avisoft, Berlin, Germany). Ultrasonic vocalizations (USVs) recordings lasted 10 min. This timeframe was selected based on our previous studies assessing the emission of USVs in dyads of unacquainted rats.^53,54^ USV recordings were converted into spectrograms by means of the software SASLab Pro 4.52 (Avisoft, Berlin, Germany) with the following settings: 512 Fast Fourier Transform (FFT)-length, Hamming window, and 75 % overlap frame set-up. Experimenters blind to the treatment groups inspected and manually cleaned all signals that could not be unambiguously classified as USVs. The SASLab Pro 4.52 software was then used to count the numbers of USVs, according to criteria previously described.^55^

### Statistical analysis

All statistical analyses were performed using GraphPad Prism® version 8 for Windows (GraphPad Software, San Diego, CA, USA) by investigators blinded to the experimental conditions. Data are expressed as mean ± SEM. The normality of data distribution was assessed using the Kolmogorov–Smirnov test for all datasets and experimental designs. When data followed a normal distribution, statistical comparisons were made using one- or two-way ANOVA followed by Tukey’s post hoc test, or unpaired Student’s *t*-test. For non-normally distributed data, the Kruskal–Wallis test followed by Dunn’s post hoc test, or the Mann–Whitney test, was applied as appropriate. Statistical significance was defined as *p* < 0.05.

## Results

### Obtainment of FL TDP-43 for *in vitro* and *in vivo* studies

#### Purification and protein quality control of FL TDP-43

human FL TDP-43 was recombinantly expressed and purified with a 6x His-tag fused to the N-terminus (see Methods for details). Final conditions of the purified protein were 50 mM 4-(2-hydroxyethyl)-1-piperazineethanesulfonic acid (HEPES), pH 8.0, 0.1% (w/v) Polyethylene Glycol 3350 (PEG 3350), 2 mM Dodecyldimethylaminoxid (LDAO), 0.25% (w/v) Octyl-β-D-Glucopyranoside (OG), 1 M NaCl. Prior to analysis, we ensured that the protein sample had a high purity, using SDS-PAGE, was properly folded, using its intrinsic fluorescence and far-UV circular dichroism signature spectra, and was dimeric and not aggregated using dynamic light scattering (DLS), as previously described.^42^

### Toxicity of FL TDP-43 *in vitro*

#### SH-SY5Y cells internalization of FL TDP-43

exogenous TDP-43 uptake was verified in SH-SY5Y cells exposed to 0.12 µM CF488A-labelled FL TDP-43 for 24 hours before fixation. Brightest green fluorescence was observed in the cytoplasmic compartments while nuclei, counterstained with Hoechst dye, appeared blue (Fig. 2 A). The protein accumulated in intracellular inclusions occasionally appearing round-shaped and thus reminiscent of liquid droplets, whereas untreated cells displayed a clear cytoplasm devoid of aggregates. These findings demonstrated that FL TDP-43 was efficiently internalized and distributed within the cytoplasm of SH-SY5Y cells, forming aggregates reminiscent of TDP-43 pathology.

**Fig. 1.**
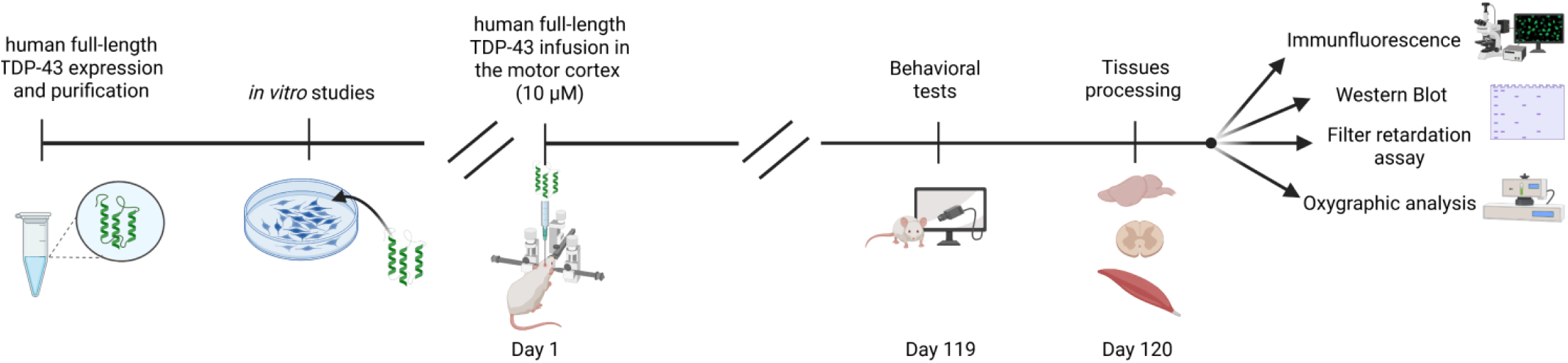
Timeline representation of the experimental workflow, outlining the sequence from protein purification to *in vivo* behavioral testing and final tissue processing.

**Fig. 2.**
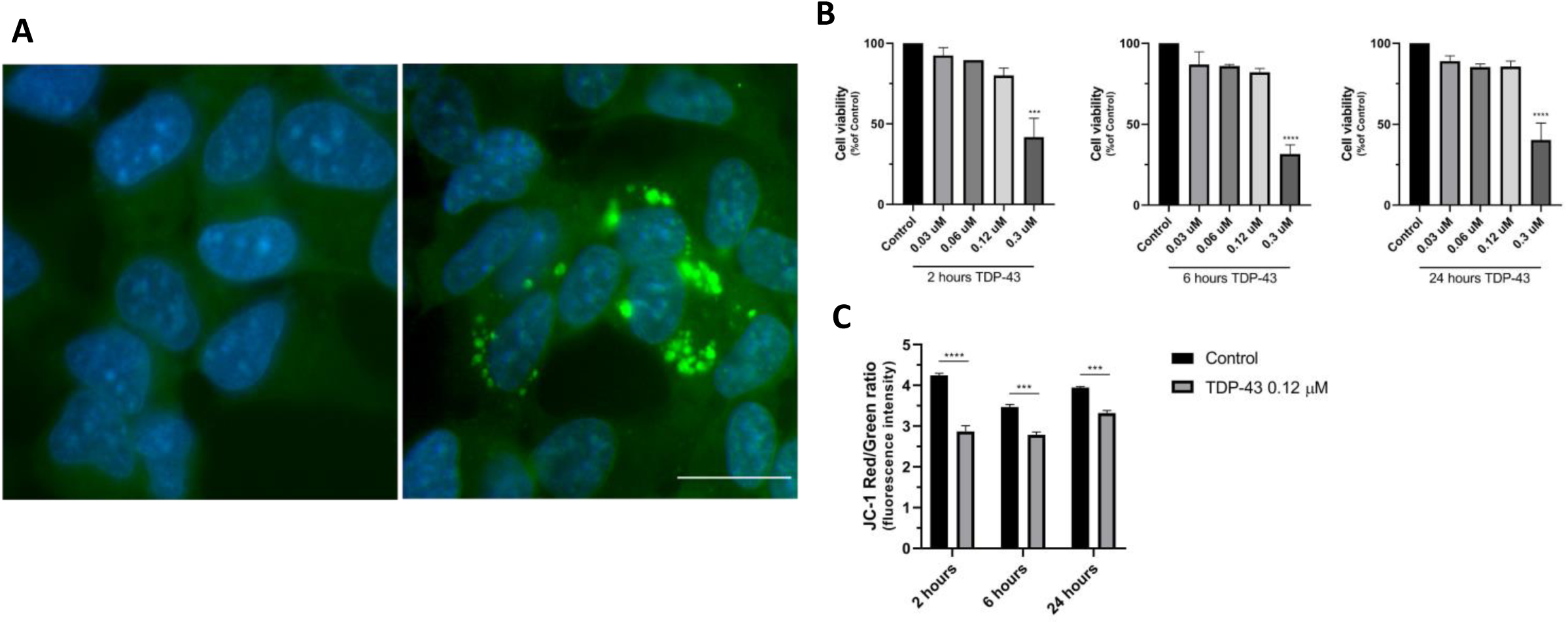
**A) TDP-43 was internalized in SH-SY5Y cells**. Protein uptake was observed after incubating for 24 hours with green fluorescent TDP-43 0.12 µM. The right panel shows TDP-43 round-shaped aggregates distributed in SH-SY5Y cytoplasm as the nuclei were blue counterstained with DAPI. Control cells (panel on the left) exhibited a cytoplasm free of aggregates. Scale bar: 20 μm. **B) TDP-43 induced cell viability loss in a concentration-dependent manner.** SH-SY5Y cells were exposed to different concentrations of TDP-43 for 2, 6, or 24 hours. Post treatment viability was measured using the MTT assay. Cell viability was expressed as a percentage of the control. Data represent three independent experiments and are presented as mean ± SEM. One-way ANOVA followed by Dunnet’s post-hoc. *** p < 0.001, **** p < 0.0001 **C) TDP-43 induced SH-SY5Y cells mitochondrial depolarization.** After treatment with 0.12 μM TDP-43 for 2, 6 and 24 hours, SH-SY5Y cells showed a reduced mitochondrial membrane potential compared to control cells as demonstrated by the decrease in JC-1 ratio between red and green emitted fluorescence. Data are expressed as fluorescence intensity. Two-way ANOVA followed by Tukey’s post-hoc. Data represent means ± SEM of N = 3 experiments. ***p < 0.001, ****p < 0.0001.

#### Cell viability of SH-SY5Y cells upon exposure to FL TDP-43

the cytotoxic effect of FL TDP-43 on SH-SY5Y cells was evaluated by the MTT assay in SH-SY5Y cells treated with increasing concentrations of the protein for 2, 6 or 24 hours. Figure 2 B shows that the highest concentration tested (0.3 µM) significantly decreased cell viability to a value of 60-70% compared to control cells. At lower concentrations (from 0.03 µM to 0.12 µM), FL TDP-43 caused a slight reduction in cell viability, although this was not statistically significant. This data indicated that FL TDP-43 exhibited the highest cytotoxic effect at 0.3 µM after 2 hours (p =0.0002), which continued up to 6 hours (p <0.0001) and 24 hours (p <0.0001).

#### Mitochondrial membrane potential in SH-SY5Y cells exposed to FL TDP-43

mitochondrial membrane potential was determined using the voltage-sensitive dye JC-1. In healthy mitochondria JC-1 accumulates in the mitochondrial matrix forming red, fluorescent aggregates due to its high concentration. In contrast, in damaged mitochondria JC-1 fails to aggregate due to poor mitochondrial enrichment caused by membrane depolarization and increased permeability, leading to loss of electrochemical potential, and maintenance of original green fluorescence. The red- and green-emitted fluorescence ratio therefore indicates the membrane polarization state, with a lower ratio indicating depolarized mitochondria as a predictor of apoptotic processes.

To assess mitochondrial potential variations over time, SH-SY5Y cells were treated with 0.12 µM FL TDP-43 for 2, 6 or 24 hours. As illustrated in Figure 2 C, TDP-43 exposure decreased JC-1 red/green fluorescence ratio, achieving its main effect after 2 hours treatment (p<0.0001). Fluorescence intensity ratio was also significantly reduced after 6 and 24 hours of cell incubation (p=0.0004 and p=0.0007, respectively), confirming the TDP-43 ability to induce long-lasting mitochondrial membrane depolarization.

### Toxicity of FL TDP-43 *in vivo* - Cortical pathology

#### Intraneuronal pTDP-43 staining

the unilateral infusion of FL TDP-43 into the rat motor cortex (Fig. 3A) led to a significant increase of pTDP-43 staining within motor cortex neurons four months post-infusion (Figs. 3B and 3 C). Specifically, One-way ANOVA revealed a significant effect among groups (F_3,20_= 2.166; p=0.0003) and Tukey’s post hoc test showed a significant increase in the ipsilateral compared to the contralateral side of TDP-43-infused brains (p=0.0007), as well as compared to both sides of vehicle-infused brains (p=0.0019) (Fig. 3B). Despite the presence of intraneuronal pTDP-43, we did not detect any neuronal loss in the same cortical region, as measured by NeuN+ cells (One-way ANOVA; F_3,16_=0,1453; p=0.5045) (Fig. 3D).

**Fig. 3.**
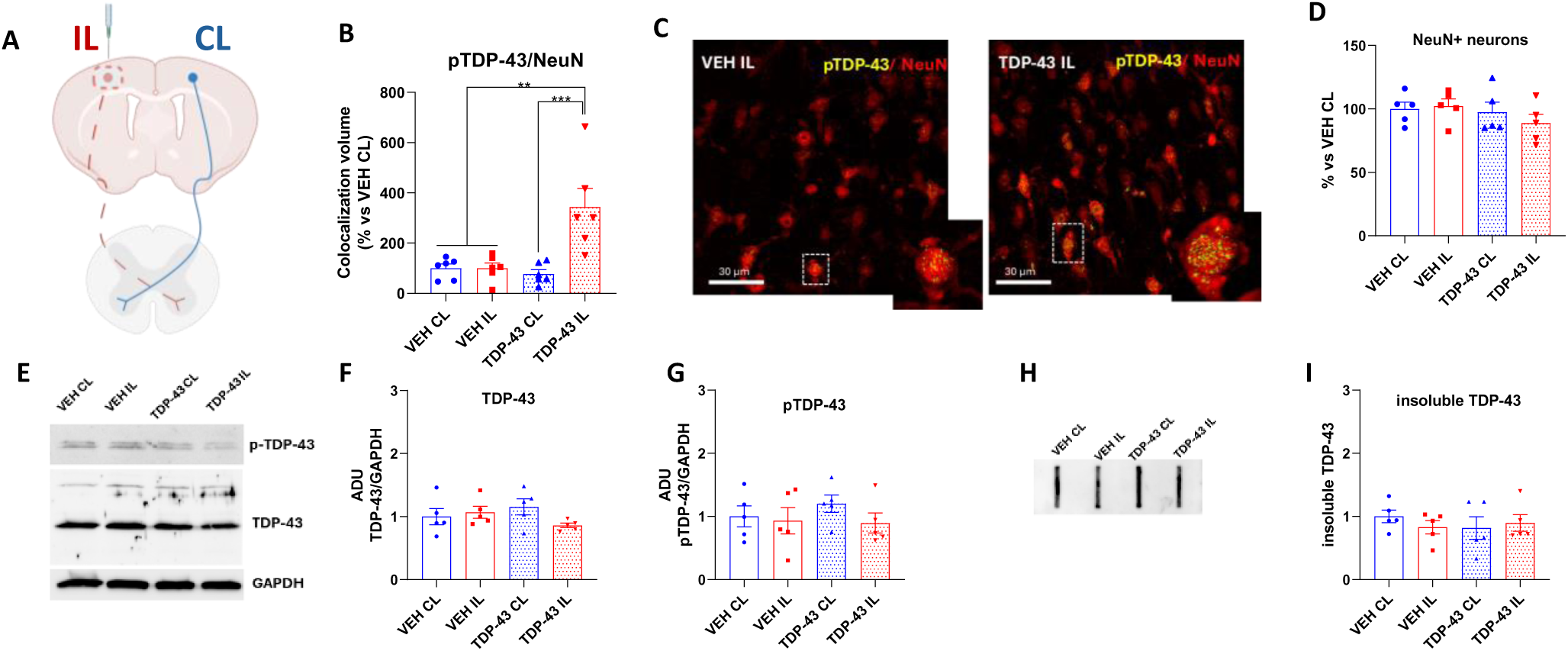
pTDP-43 aggregates in the motor cortex without affecting neuronal viability. **A)** Schematic representation of the intracortical infusion side of FL TDP-43 (IL, ipsilateral, dashed red line) and vehicle (CL, contralateral, blue line). **B)** Total volume occupied by pTDP-43 colocalized with NeuN+ cells in the motor cortex four-months post-infusion. Values represent the mean ± SEM (One-way ANOVA followed by Tukey’s post hoc test.) ***p<0.001 vs TDP-43 IL; **p<0.01 vs TDP-43 IL. **C)** Confocal images showing pTDP-43 (yellow) in NeuN+ cells (red). Scale bar: 30 μm. Dashed red square in the picture correspond to the affected motor cortex site (ipsilateral to the injection side). **D)** Number of NeuN+ neurons measured within the motor cortex. Values represent the mean ± SEM (One-way ANOVA followed by Tukey’s post hoc test). **E-I**) Representative western blot analysis (**E**) and relative quantifications of the full-length TDP-43 (**F**) and pTDP-43 (**G**). The graphs represent the densitometric analyses of TDP-43 and phospho-TDP-43 levels normalized using GAPDH as loading control. Data are expressed as arbitrary densitometric units (ADU). Each bar represents the mean ± SEM of five independent replicates (ANOVA with Tukey’s post hoc test among groups). **H-I**) Representative filter retardation assay (**H**) and relative densitometric analysis (**I**) of C-terminal insoluble TDP-43 species. Value represents the mean ± SEM of five independent replicates (ANOVA with Tukey’s post hoc test among groups). Please note that dotted red columns in the graphs always refer to the TDP-43-affected side (i.e., ipsilateral to the injection side for the motor cortex and contralateral to the injection side for the spinal cord and muscle). VEH = vehicle.

We then evaluated both TDP-43 and pTDP-43 protein levels in the whole motor cortex from a separate cohort of rats TDP-43-infused and vehicle-infused as described above, by Western blot analysis (Fig. 3D-F). Densitometric analyses showed no statistically significant differences among the samples (One-way ANOVA: TDP-43, F_3,16_=1,1443; p= 0.2673; pTDP-43, F_3,16_=0,6504; p= 0.5941). Finally, we investigated the possible formation of insoluble high–molecular-weight TDP-43 species using filter retardation assay (Fig 3G-H). Also in this case, no differences were observed (One-way ANOVA: F_3,16_=0,4012; p=0.7541), suggesting that the bulk nature of these assays, which measure proteins within the whole tissue lysate, may have diluted signals from neuronal inclusions.

### Integrity of protein quality control pathways

motor cortex extracts from TDP-43-infused and vehicle-infused rats, showed no changes in the expression levels of protein quality control (PQC) system key markers (i.e. the chaperones and co-chaperones HSPAs, HSPB8 and BAG3 proteins, the autophagy receptor SQSTM1/p62 and the autophagy marker MAP1LC3B), known to participate in the degradation of insoluble toxic TDP-43 species (Fig. S1).

### Mitochondrial damage

the intracortical infusion of FL TDP-43 reduced the staining for the fourth subunit of cytochrome c oxidase CIV within NeuN+ cells in the motor cortex. Specifically, the Kruskal-Wallis test revealed a significant effect among groups (p=0.0064) and Dunn’s post hoc test showed a significant reduction in CIV expression in the ipsilateral side of TDP-43-infused brains compared to both the contralateral and ipsilateral sides of the vehicle-infused brains (p=0.0258 and p=0.0368, respectively) (Fig. 4A). Interestingly, a trend to reduction was also observed in the contralateral motor cortex of TDP-43 infused rats (Fig. 4A).

**Fig. 4.**
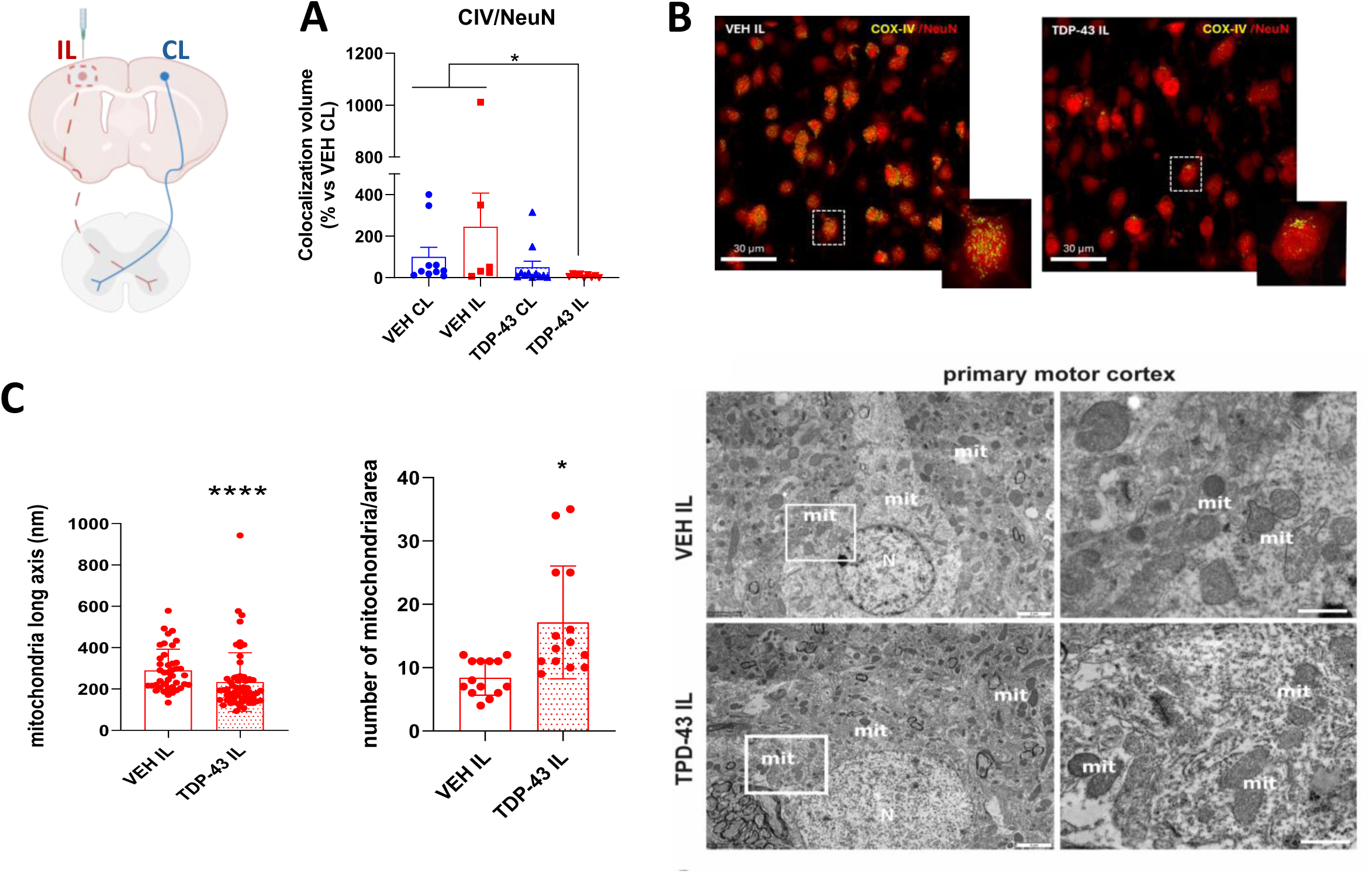
TDP-43 infusion induces alteration in mitochondrial activity and morphology in the motor cortex. **A)** Total volume occupied by CIV colocalized with NeuN+ cells in the motor cortex. Values represent the mean ± SEM (Kruskall-Wallis followed by Dunn’s post hoc test) *p<0.05 vs TDP-43 IL. **B)** Confocal images showing COX-IV staining (yellow) in NeuN+ cells (red). Scale bar: 30 μm. Dashed red square in the picture correspond to the affected motor cortex site (ipsilateral to the injection side). **C)** Histograms and relative images showing the mitochondria length (major axis) and number in the motor cortex as measured by TEM analysis. Values represent the mean ± SEM (Mann-Whitney test) ****p<0.0001; *p<0.05. Please note that dotted red columns in the graphs always refer to the TDP-43-affected side (i.e., ipsilateral to the injection side for the motor cortex and contralateral to the injection side for the spinal cord and muscle). C = complex; CL = contralateral to the injection site; IL = ipsilateral to the injection site; VEH = vehicle.

The investigation of dataset showed a non-normal distribution, suggesting a high variability in the content of CIV within cortical neurons. To further investigate this aspect, data was expressed as a frequency distribution of CIV volume within NeuN+ cells. Frequency distribution analysis revealed two main cell populations for each treatment (i.e., displaying low CIV labelling and high CIV labelling). Based on this data distribution, a cut-off was set and groups were statistically compared (Fig. S2). One-way ANOVA revealed a significant effect among groups (F_3, 18_ = 7,678, p=0.0016) and Tukey’s post hoc showed a significant difference for each population considered, revealing an increase percentage of cells with low CIV labelling, and a decreased percentage of cells with high labelling, in the motor cortex ipsilateral to TDP-43 infusion (Fig. S2).

TEM analysis revealed a marked remodeling of mitochondrial morphology in the ipsilateral motor cortex following FL TDP-43 infusion. In control conditions, neuronal mitochondria displayed the expected elongated profile with well-preserved cristae and a homogeneous matrix. In contrast, on the TDP-43–affected side (ipsilateral), mitochondria appeared significantly shorter and more fragmented, with a clear shift toward small, rounded profiles suggestive of increased fission and/or reduced fusion. Quantification confirmed a robust reduction in mitochondrial length (major axis) in ipsilateral samples compared to the contralateral hemisphere (Fig. 4C; p<0.0001), together with a modest but significant increase in the total number of mitochondrial profiles per field (p=0.0273).

#### *in vivo* toxicity of FL TDP-43 - Spinal cord pathology

##### Bilateral pathological spreading

the intracortical infusion of FL TDP-43 caused an increase of pTDP-43 labelling within the contralateral cervical spinal cord, which is anatomically connected with the TDP-43 infused cortex (see scheme in Fig. 5A showing pyramidal decussation of nerve fibers) (Fig. 5B-C). Kruskal Wallis test revealed a significant effect among groups, and Dunn’s post hoc test showed a significant increase of pTDP-43 in the spinal cord contralateral to the TDP-43 intracortical infusion site; in addition, a significant increase was also observed in the contralateral spinal cord of intracortically vehicle-infused rats compared to the ipsilateral side (p=0.0163 and p=0.03, respectively) (Figs. 5C).

**Fig. 5.**
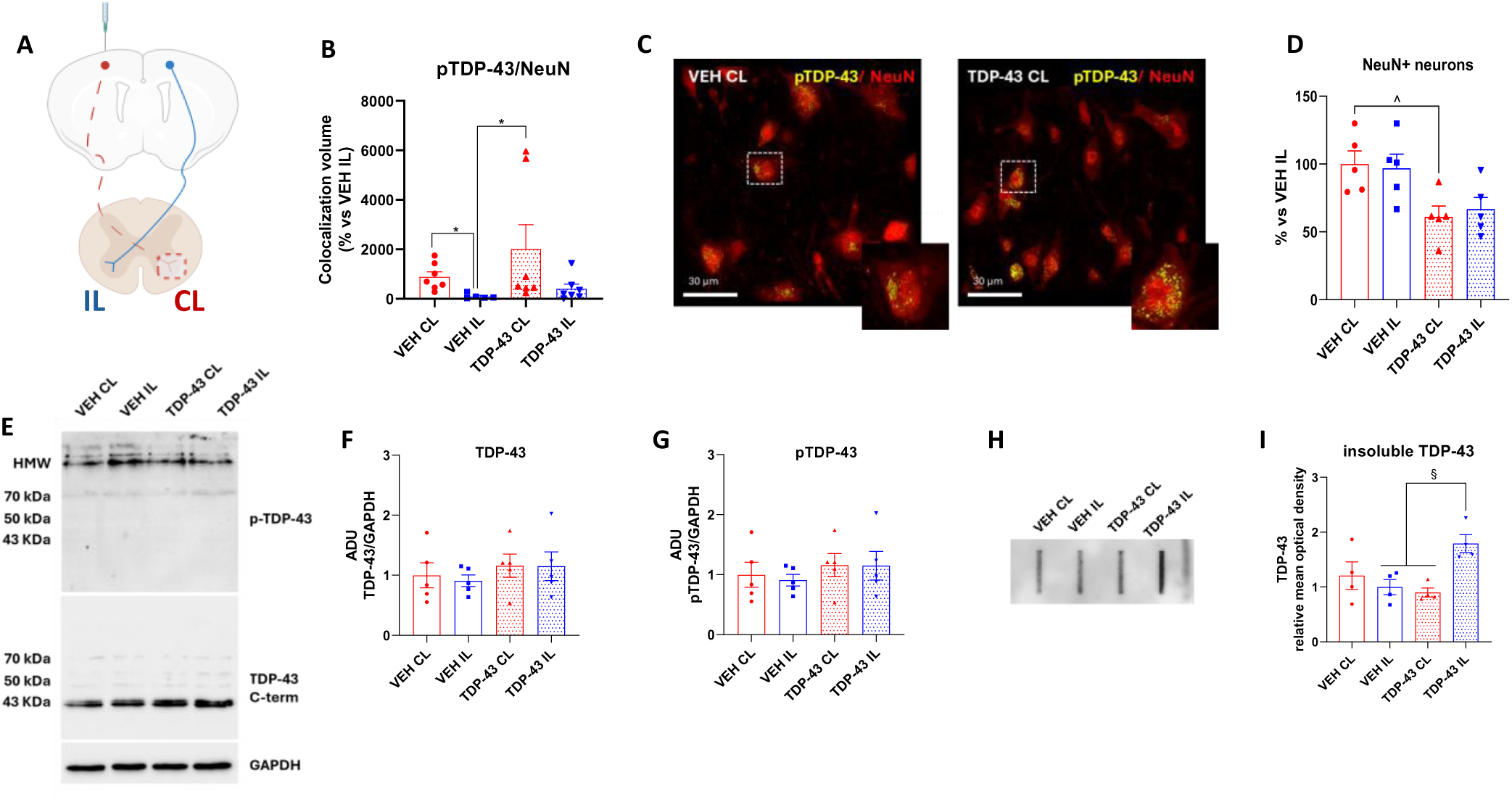
pTDP-43 aggregates in the cervical tract of the spinal cord and induces neuronal degeneration. **A)** Schematic representation of the intracortical infusion side of FL TDP-43 (IL, ipsilateral, dashed red line) and vehicle (CL, contralateral, blue line). **B)** Number of NeuN+ neurons measured within the cervical tract of the spinal cord. Values represent the mean ± SEM (One-way ANOVA followed by Tukey’s test) ^p<0.05 vs TDP-43 CL. **C)** Total volume occupied by pTDP-43 colocalized with NeuN+ cells in the motor cortex four-months post-infusion. Values represent the mean ± SEM (Kruskall-Wallis followed by Dunn’s post hoc test.) *p<0.05 vs TDP-43 CL and VEH CL. **D)** Confocal images showing pTDP-43 (yellow) in NeuN+ cells (red). Scale bar: 30 μm. Dashed red square in the picture correspond to the affected spinal cord site (contralateral to the injection side). **E-G**) Representative western blot analysis (**E**) and relative quantifications of the C-terminal TDP-43 (**F**) and pTDP-43 (**G**) species. The graphs represent the densitometric analyses of C-terminal and phospho-TDP-43 signals normalized using GAPDH as loading control. Data are expressed as arbitrary densitometric units (ADU). Each bar represents the mean ± SEM of five independent replicates (ANOVA with Tukey’s post hoc test among groups). **H-I**) Representative filter retardation assay (**H**) and relative densitometric analysis (**I**) of C-terminal insoluble TDP-43 species. Value represents the mean ± SEM of five independent replicates (ANOVA with Tukey’s post hoc test among groups). §p<0.05 vs TDP-43 IL. Please note that dotted red columns in the graphs always refer to the TDP-43-affected side (i.e., ipsilateral to the injection side for the motor cortex and contralateral to the injection side for the spinal cord and muscle). CL = contralateral to the injection site; IL = ipsilateral to the injection site; VEH = vehicle.

In the cervical spinal cord, the intracortical infusion of FL TDP-43 also induced a significant neuronal loss in the side contralateral to infusion (Fig 5D). One-way ANOVA revealed a significant effect among groups (F_3,16_=0,2291; p=0.0157) and Tukey’s post hoc test showed a significant reduction of NeuN+ cells in the contralateral spinal cord of TDP-43-infused rats as compared to the same region from vehicle-infused rats (p=0.0407) (Fig. 5D). In addition, a reduction of NeuN+ cells, although not statistically significant, was observed in the ipsilateral spinal cord of TDP-43-infused rats (Fig. 5D).

Western blot analyses of whole cervical spinal cord extracts and their relative quantifications (Fig. 5E-G) revealed small but non-statistically significant increases in the levels of either total TDP-43 or pTDP-43 form among the samples (One-way ANOVA: TDP-43, F_3,16_=1,1443; p= 0.2673; pTDP-43, F_3,16_=0,1348; p=0.9379). However, filter retardation assay analysis (Fig. 4H-I) showed a significant increase in insoluble high–molecular-weight TDP-43 species in the spinal cord samples ipsilateral to the infusion site of TDP-43-infused rats compared to both samples contralateral to the infusion site of TDP-43-infused rats and those ipsilateral to the infusion site of vehicle-infused rats (One-way ANOVA: F_3,12_=5,541; p=0.0127; Tukey’s post hoc: p=0.0135 and p=0.0279, respectively). This accumulation of insoluble forms warrants further investigation to clarify its underlying mechanisms and potential implications. As previously observed for the motor cortex, total protein extracts from the cervical spinal cord tracts after FL TDP-43 intracortical infusion did not reveal any statistically significant changes in the protein levels of key PQC markers analyzed (One-way ANOVA: HSPAs F_3,15_=1,793; p=0.1919; HSPB8: F_3,16_=0,7551; p=0.5354; p62 F_3,16_=0,2397; p= 0.8673; LC3B F_3,16_=0,0640; p=0.9781) (Fig. S3).

### Mitochondrial damage

intracortical infusion of the FL TDP-43 led to a reduced expression of CIV within NeuN+ cells in the cervical spinal cord. Kruskal-Wallis test revealed a significant effect among groups and Dunn’s post hoc test showed a significant reduction in the expression of CIV in the spinal cord contralateral to the infusion site of TDP-43-infused rats, compared to vehicle-infused animals (p=0.0483) (Fig. 6A). Moreover, a trend to reduction was also observed in the ipsilateral spinal cord of TDP-43 infused rats (Fig. 6A).

**Fig. 6.**
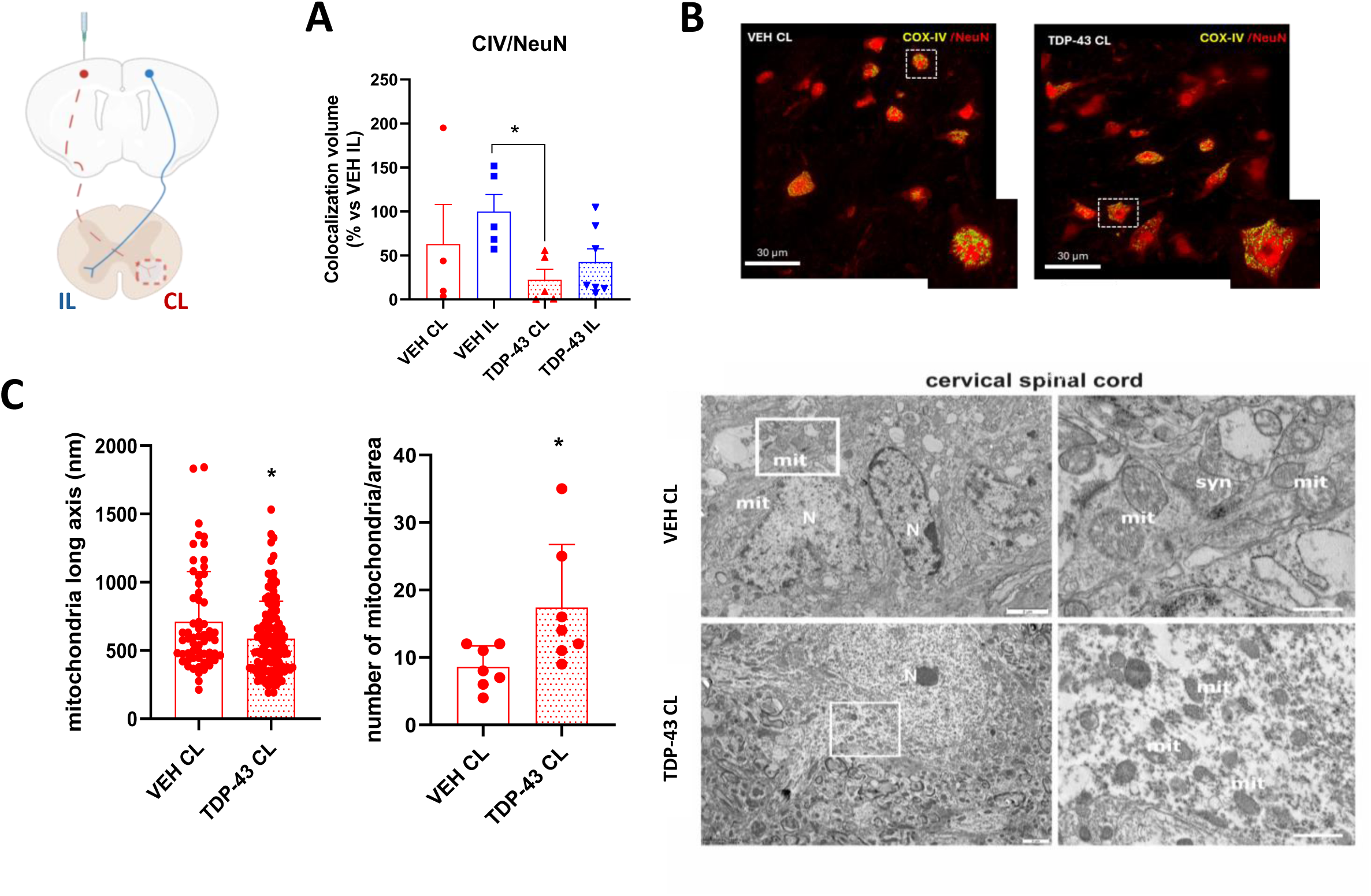
TDP-43 infusion induced alteration in mitochondrial activity and morphology in the spinal cord. **A)** Total volume occupied by CIV colocalized with NeuN+ cells in the spinal cord. Values represent the mean ± SEM (Kruskall-Wallis followed by Dunn’s post hoc test) *p<0.05 vs TDP-43 CL. **B)** Confocal images showing CIV staining (yellow) in NeuN+ cells (red). Dashed red square in the picture correspond to the affected spinal cord site (contralateral to the injection side). **C)** Histograms and relative images showing the mitochondria length (major axis) and number in the cervical tract of the spinal cord as measured by TEM analysis. Values represent the mean ± SEM (Mann-Whitney test) *p<0.05. Please note that dotted red columns in the graphs always refer to the TDP-43-affected side (i.e., ipsilateral to the injection side for the motor cortex and contralateral to the injection side for the spinal cord and muscle). C = complex; CL = contralateral to the injection site; IL = ipsilateral to the injection site; VEH = vehicle.

TEM analysis revealed a distinct but convergent pattern of mitochondrial impairment in the cervical spinal cord after FL TDP-43 intracortical infusion. Consistent with the confocal quantification, which revealed a significant reduction of CIV–positive volume in NeuN+ neurons on the TDP-43–affected side (Fig. 6A–B; p=0.0347), TEM analysis showed clear signs of altered mitochondrial dynamics in spinal motor neurons. Ultrastructurally, mitochondria in the contralateral hemicord of TDP-43-infused rats appeared less elongated and more heterogeneous compared with the contralateral side of vehicle-infused rats. While the overall matrix density remained relatively preserved, the mitochondrial profiles were noticeably shorter, with fewer instances of the long tubular organelles typically observed in healthy motor neurons. Quantitative morphometry confirmed a significant reduction in mitochondrial length on the affected side (Fig. 6C; p=0.0347), accompanied by a mild but detectable increase in the number of mitochondrial profiles per field, suggestive of a shift toward enhanced fission or defective fusion.

### *in vivo* toxicity of FL TDP-43 - Skeletal muscle pathology

Four months after FL TDP-43 infusion into the primary motor cortex, significant bilateral alterations of skeletal muscle bioenergetics were found (Fig 7A-D). Specifically, bioenergetics of isolated mitochondria from the contralateral gastrocnemius showed a significant increase in oxidative capacity at CII (p=0.03, Fig. 7 B). Uncoupled respiration in contralateral CII was also significantly increased in TDP-43 samples with a significance level at p=0.037 (Table 1 Suppl. Material).

**Fig. 7.**
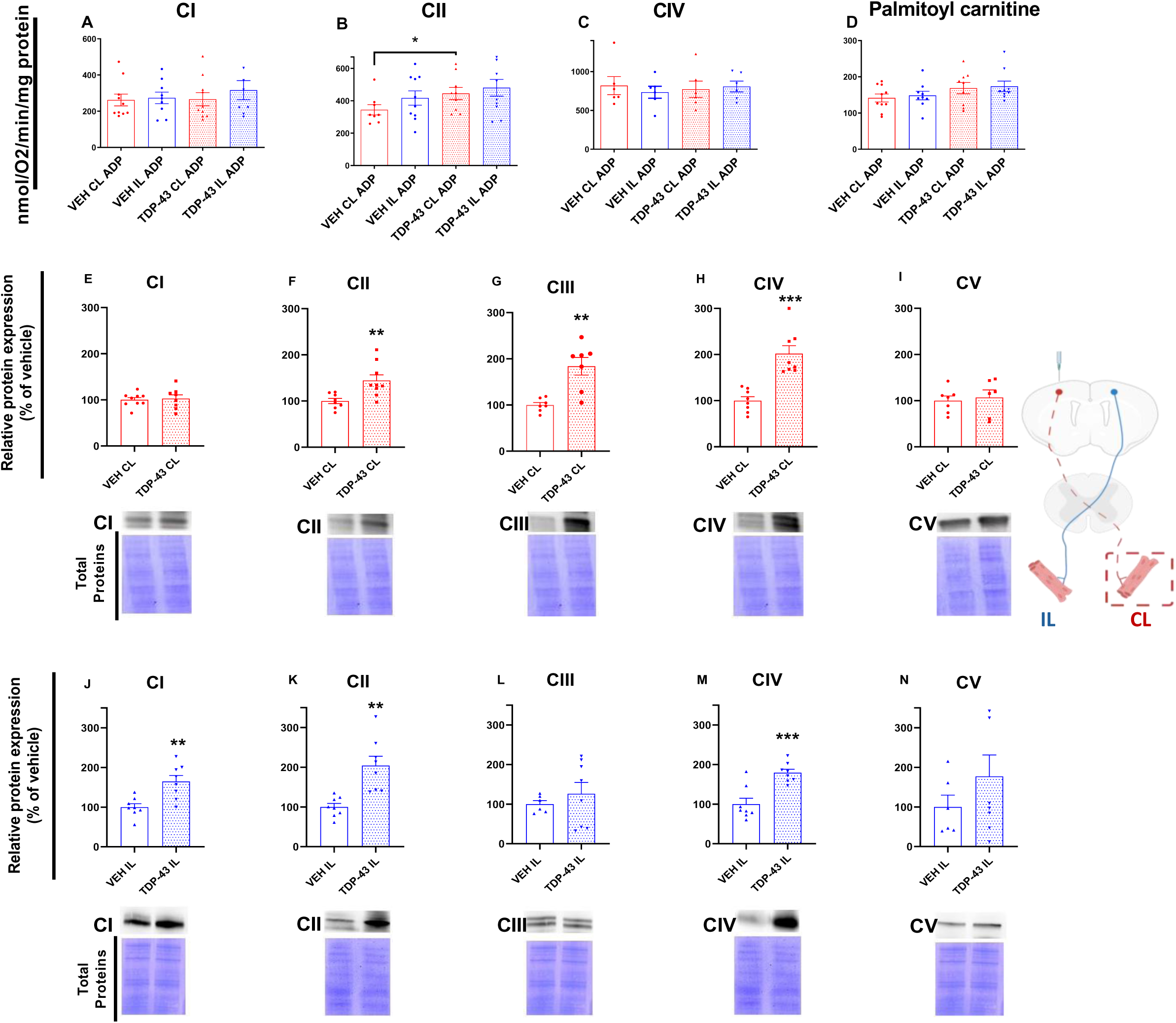
Bioenergetics on skeletal muscle mitochondria isolated from CL or IL gastrocnemius. **(A-D).** Data show oxygen consumption following ADP or uncoupler administration in mitochondria of vehicle or TDP-43-treated animals. Most of the values show a tendency to be higher in TDP-43-treated animals, but they reached statistical significance only in CL CII (B) (p<0.05 Mann-Whitney test). Dashed red square in the picture correspond to the affected muscle (gastrocnemius) (contralateral to the injection side). Western blot on electron transport chain complexes expression in isolated mitochondria from CL or IL gastrocnemius (E-N). After TDP-43 treatment several complexes increased their expression. Data are normalized for total protein and expressed as a percentage of vehicle. ** p<0.001, *** p<0.0001, Student’s t-test. Please note that dotted red columns in the graphs always refer to the TDP-43-affected side (i.e., ipsilateral to the injection side for the motor cortex and contralateral to the injection side for the spinal cord and muscle). C = complex; CL = contralateral to the injection site; IL = ipsilateral to the injection site; VEH = vehicle.

We also evaluated protein expression of markers for mitochondrial complexes (CI, NDUFB8; CII, SDHB; CIII, UQCRC2; CIV, MTCO1; CV, ATP5A) by western blot run in mitochondria isolated from the contralateral and ipsilateral gastrocnemius (Figs. 7E-N). Densitometry results showed that in the contralateral gastrocnemius, the expression of CII, III, and IV increased after FL TDP-43 intracortical infusion. In the ipsilateral gastrocnemius, the increase was statistically significant for CI, II, and IV, while the expression of ipsilateral III and V showed no significant increase. Indeed, statistical analysis showed significant values with p =0.0062 and p=0.0011 for contralateral CII and III, respectively, and p=0.0024 and p=0.0010 for ipsilateral CI and II, and, last, statistically significant values with p=0.001 and p=0.0004 for CIV in both contralateral and ipsilateral gastrocnemii, respectively.

These data confirmed that the intracortical FL TDP-43 infusion affected bilaterally skeletal muscle mitochondria. Changes in the expression of single proteins in complexes were only partially transduced into a variation of OXPHOS, in line with the efficient functional reserve capacity of mitochondria and the complex interaction of proteins inside the complexes.

Finally, to determine whether FL TDP-43 infusion induced any changes in the expression levels, phosphorylation, or aggregation of endogenous TDP-43, as well as of PQC markers in rat muscle, western blot and filter retardation assay analyses were performed on protein extracts from gastrocnemius muscles of the different experimental groups. Densitometric analyses showed no differences among the groups for any of the proteins analyzed (One-way ANOVA: TDP-43 F_3,16_= 0,7067, p=0,5619; pTDP-43 F_3,12_=0,6403, p=0,6036; insoluble TDP-43 F_3,16_=0,03959, p=0,9891; HSPAs F_3,16_=0,04178, p=0,9882; HSPB8: F_3,16_=0,1645, p=0,9187; p62 F_3,16_=0,7194, p=0,5548; LC3B-I F_3,16_=2,691, p=0,0811; LC3B-II F_3,16_=2,638, p=0,0851) (Figs. S4 and S5).

### *in vivo* toxicity of FL TDP-43 - Behavioral phenotypes

#### Sensorimotor functions, muscular strength and fatigue

to assess the impact of brain FL TDP-43 infusion on sensorimotor function, rats were subjected to the challenging beam walking test. FL TDP-43 infusion resulted in significant sensorimotor deficits on the 15 mm and 10 mm beams. Specifically, on the 15 mm beam, t-test revealed a significant increase in the total number of errors per step made with total (i.e., combined ipsilateral and contralateral) paws of TDP-43-infused rats compared to controls (t = 3.061, df = 18, p=0.0067) (Fig. 8A). Further analysis showed a significant increase in the number of errors per step made with both the ipsilateral (t = 2.783, df = 18, p=0.0123) (Fig. 8A) and contralateral paws (t = 2.751, df = 18, p=0.0132) (Fig. 8A). On the 10 mm beam, FL TDP-43 infusion resulted in a significant increase in the number of errors per step for the contralateral paws only (t = 2.250, df = 18, p=0.0372) (Fig. 8A), while a trend toward increased errors was observed for both the total and ipsilateral paws.

**Fig. 8.**
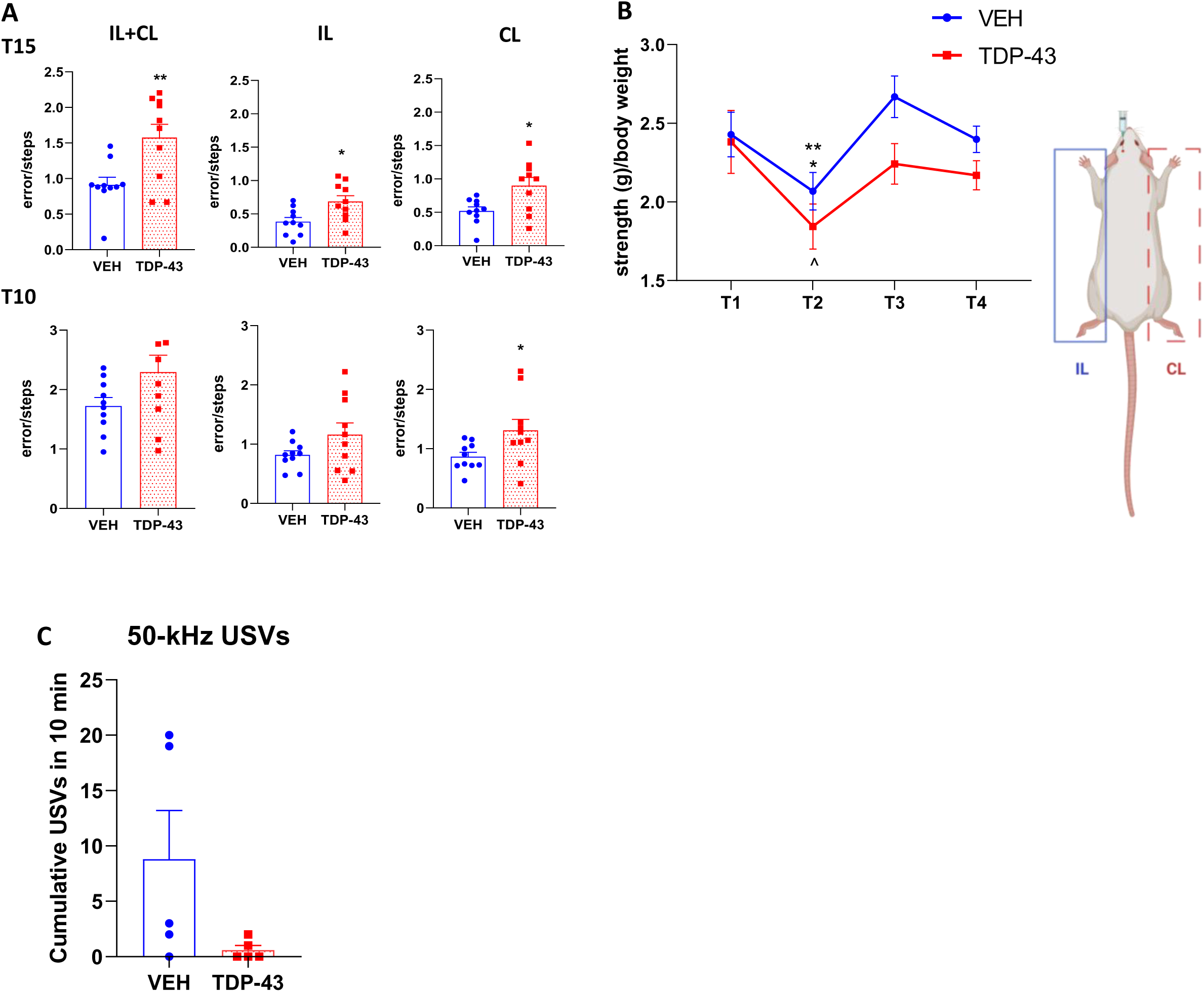
TDP-43-treated rats displayed sensorimotor deficits and increased fatigue, with a trend towards decreased 50-kHz vocalizations. **A)** Sensorimotor deficits were evaluated by the challenging beam walk test. T15 and T10 correspond to the 15 mm and 10 mm beam, respectively. Graphs represent ipsilateral and contralateral paws respect to the injection site (left), ipsilateral paws respect to the injection site (center) and contralateral paws respect to the injection site (right). Values represent the mean ± SEM (t-test) **p<0.01; *p<0.05. **B)** Muscular strength and fatigue were evaluated by grip strength test. Values represent the mean ± SEM (Two-way ANOVA followed by Tukey’s post hoc test) **p<0.01 vs VEH T3; *p<0.05 vs VEH T4; ^p<0.05 vs TDP-43 T1. **C)** The figure reveals the total number of 50-kHz USVs emitted by dyads of rats. Values represent the mean ± SEM (Mann-Whitney test). N=5 dyads for VEH-infused rats; N=5 dyads for TDP-43-infused rats CL = contralateral to the injection site; IL = ipsilateral to the injection site; USVs=ultrasonic vocalizations; VEH = vehicle.

The grip test was used to assess muscular strength and fatigue. Two-way ANOVA revealed a significant effect of time (F_3,52_ = 11,20; p<0.0001) but neither a significant effect of treatment (F_1,18_ = 2,525; p=0.1295) nor a significant interaction time x treatment (F_3,52_ = 1,339; p=0.2719) (Fig. 8B). A trend towards reduced muscular strength was observed in TDP-43-infused rats compared to the vehicle-treated-rats for the time points T2, T3, and T4, corresponding to 1 minute (T2) and 3 hours (T3 and T4) after the first trial (T1) (Fig. 8B).

#### Emission of 50-kHz ultrasonic vocalizations

dyads of TDP-43-infused rats showed a trend towards an increased emission of 50-kHz USVs when placed inside the test cage compared with dyads of vehicle-infused rats (Fig. 8C). Nevertheless, this effect did not reach statistical significance, as shown by Mann Whitney test (U=4, sum of ranks= 36, 19, p=0.1032). No emission of 22-kHz USVs was recorded in the present study.

## Discussion

TDP-43 is a ubiquitously expressed RNA/DNA-binding protein essential for RNA metabolism and neuronal homeostasis.^56^ In ALS and FTLD, hyperphosphorylated TDP-43 is a major component of cytoplasmic inclusions in cortical and spinal motor neurons, representing a defining pathological hallmark.^10,57^

Increasing evidence indicate that TDP-43 can be released into the extracellular environment within vesicles or as a free protein, with increased release under pathological conditions.^14,58–60^ This suggests that extracellular TDP-43 may propagate in a prion-like manner contributing to disease propagation. However, while the seeding and spreading of patient-derived TDP-43 aggregates have been demonstrated in cultured cells and in a single study *in vivo*,^36,59,61^ the intrinsic pathogenic potential of native FL TDP-43 *in vivo* has remained largely unexplored.^39,40,62^

In the present study, we provide direct evidence that exogenous, native FL TDP-43 is sufficient to induce toxicity and drive pathological spreading *in vivo*. We first show that purified human FL TDP-43 was efficiently internalized by neuronal cells *in vitro*, where it induced aggregation, impaired mitochondrial function, and reduced cell viability.^41,63^ Extending these findings *in vivo*, intracortical infusion of FL TDP-43 triggered a progressive and anatomically defined centrifugal propagation of pathology along the corticospinal axis.^41^ This spreading was defined by intraneuronal pTDP-43 inclusions, accumulation of high-molecular-weight species, neuronal dysfunction, and region-specific neurodegeneration. TDP-43 toxicity ultimately disrupted peripheral muscle bioenergetics and function, resulting in a behavioral phenotype featuring key aspects of human symptomatology, such as motor impairment and increase in muscular fatigue.

### CNS toxicity and propagation

Exogenous FL TDP-43 induced widespread CNS toxicity. Notably, pathology propagated corticofugally from the motor cortex to the spinal cord, consistent with previous reports identifying pTDP-43 accumulation in the CNS as a pathological hallmark of ALS.^64^ However, the burden of pathology was region-specific. In the motor cortex, pTDP-43 accumulation was not associated with overt neuronal loss but coincided with marked mitochondrial alterations, suggesting early neuronal dysfunction.^64,65^ In contrast, the spinal cord contralateral to the protein infusion side, displayed pTDP-43 inclusions and a significant motor neuron degeneration, particularly affecting large ventral horn neurons, indicating increased vulnerability at distal sites.^66,67^ Importantly, we show that toxicity spread beyond regions with overt pTDP-43 deposition in the spinal cord. Despite limited detection of pTDP-43 inclusions, partial neuronal loss and mitochondria disruption were observed in the ipsilateral spinal cord of TDP-43-infused rats, together with high levels of insoluble TDP-43 species, as shown by the filter retardation assay. This assay captures all detergent-insoluble TDP-43 species regardless of phosphorylation, suggesting that early accumulation of non-phosphorylated, detergent-insoluble TDP-43 may precede overt inclusion formation and contribute to neuronal damage. In contrast, WB did not fully recapitulate the immunofluorescence findings, likely due to the bulk nature of this assay, which dilutes signals from neuronal inclusions across the whole tissue lysate. Notably, pTDP-43 pathology increasingly affects non-neuronal cells along ALS progression, which may align with an early pathological phase of our model. ^64,72,73^ Consistently, no changes were detected in the expression of PQC components in either cortex or spinal cord, which are typically altered only at later stages in ALS models.^74,75^

Interestingly, pTDP-43 inclusions were also detected in the spinal cord of vehicle-treated rats, suggesting that the infusion procedure itself may induce local pTDP-43 accumulation as part of a stress response without causing significant motor neuron loss.^68–71^

Together, these findings indicate that FL TDP-43 toxicity propagates along the corticospinal axis and exerts deleterious effects even before classical pathological hallmarks become fully detectable.

### Mitochondrial disfunction as an early driver of pathology

A key finding of this study is the early and widespread mitochondrial vulnerability induced by FL TDP-43. In line with previous findings *in vitro,* FL TDP-43 reduced mitochondrial membrane potential of SH-5YSY cells, even at concentrations that did not affect cell viability, indicating that mitochondrial dysfunction preceded overt cell death.^80^

*In vivo*, cortical infusion of FL TDP-43 led to mitochondrial alterations in both cortical and spinal neurons, indicated by reduced expression of mitochondrial respiratory chain components and profound ultrastructural remodeling, including mitochondrial fragmentation. The concordance between confocal and ultrastructural analyses revealed not only changes in mitochondrial markers but also a remodeling of mitochondrial architecture in vulnerable neurons.^76^ Although CIV activity in the CNS was not directly measured, its expression is widely regarded as a reliable indicator of mitochondrial functional integrity.^83,84^ Therefore, the increased abundance of shorter mitochondria likely reflects mitochondrial fragmentation rather than enhanced biogenesis. These findings reinforce the concept that mitochondria are central mediators of TDP-43 toxicity and identify their dysfunction as an early pathogenic event triggered by native FL TDP-43. Results align with previous reports in WT and mutant TDP-43 models, showing mitochondrial vulnerability to mislocalized and aggregated proteins,^29,77–79,81,82^ and with a recent study showing that CIV deficiency in rats recapitulate features of ALS pathology.^86^

Overall, these findings suggest that cortical FL TDP-43 infusion induces mitochondrial alterations across the CNS, representing an early mechanism of dysfunction relevant to ALS pathology.

Within the framework of pathological biomarkers, our findings suggest that early TDP-43–induced alterations could support prodromal diagnosis, complementing established markers such as neurofilaments that primarily reflect overt neurodegeneration.^91,92^ In line with this concept, we recently identified a translational TDP-43–induced transcriptomic–miRNA signature in the rat model used here, detected across CNS regions and peripheral blood cells. In patients, the same signature was also validated in peripheral blood cells from ALS but not from PD, supporting disease-specific peripheral molecular changes as early indicators of TDP-43 pathology (Manchinu et al., under revision).

### Peripheral propagation and muscle involvement

Notably, we demonstrate that FL TDP-43 toxicity extends beyond the CNS to peripheral tissues, reaching skeletal muscle. In gastrocnemius muscle, mitochondrial alterations were evident, including bilateral dysregulation of respiratory chain components and altered bioenergetic function, supporting the peripheral spreading of TDP-43 pathology. This upregulation did not fully correspond to a bilateral alteration of OXPHOS function, as assessed by oxygraphic analysis. While increased expression of Cs II–IV was observed, this did not translate into a proportional increase in respiratory activity, which was detected only for contralateral CII. This discrepancy likely reflects a compensatory response, whereby increased expression of mitochondrial complex subunits does not necessarily result in fully functional complexes. Consistent with these findings, early-stage ALS patients show increased CII activity with preserved C IV function, while a decline in C IV is observed at later disease stages.^86–88^

Despite these bioenergetic alterations, no increase in pTDP-43 or C-terminal fragments was detected in skeletal muscle by WB. This may reflect a more efficient PQC system in muscle relative to motor neurons, facilitating the clearance of misfolded TDP-43. Consistently, skeletal muscle displays higher proteasomal and autophagic activity, with PQC components generally upregulated only at later stages of disease.^74,89^ In line with this, no significant changes in PQC markers were detected in skeletal muscle, including chaperones (HSPAs, HSPB8, BAG3) and autophagy-related proteins (SQSTM1/p62, MAP1LC3B), supporting an early-stage pathological context.

Notably, pTDP-43 accumulation has been described in skeletal muscles of ALS patients prior to the onset of clinical symptoms and electromyographic alterations, highlighting a limitation of the present study.^90^

Together, these findings provide evidence for a brain-to-muscle axis of TDP-43 toxicity, linking central pathology to peripheral dysfunction.

### Behavioral correlates of FL TDP-43 toxicity

The cellular and molecular alterations induced by FL TDP-43 were accompanied by functional deficits. TDP-43-infused rats exhibited impaired motor coordination, increased stepping errors, and enhanced fatigue, despite preserved maximal strength. These features are consistent with early-stage ALS and align with the observed mitochondrial dysfunction and spinal cord pathology. Motor performance was evaluated using the challenging beam test, a well-established measure of motor coordination and balance in rodents.^93^ TDP-43-infused rats exhibited a bilateral increase in stepping errors on the wider beam, indicative of subtle motor impairment at an early disease stage, and consistent with the bilateral pathological alterations identified in the spinal cord and muscle. In line with this result, studies in rats with CIV deficiency indicate that more pronounced motor deficits typically emerge at later stages of disease.^85^ Muscle strength was evaluated using the grip strength test, a common measure of early functional impairment in ALS.^49–52^ While baseline strength was comparable between groups, TDP-43-infused rats showed a marked reduction in force one minute after the first trial, with no subsequent recovery, indicating increased fatigability despite preserved maximal strength. This suggests that fatigue is an early feature of the model, associated with mild motor dysfunction. Results align with clinical observations, where fatigue in ALS can occur independently of overt muscle weakness and shows limited correlation with maximal strength.^94^ The absence of clear strength deficits is therefore consistent with an early disease stage and may reflect compensatory contributions from more resistant motor neuron populations.^95,96^

TDP-43-infused rats exhibited a trend toward reduced emission of 50-kHz USVs during dyadic interactions, a paradigm typically associated with positive affective states.^53,54,97^ This reduction may reflect early alterations in emotional processing, as reported in non-demented ALS patients.^98,99^ However, given that USV production relies on the coordinated activity of laryngeal and respiratory muscles innervated by cervical motor neurons, a contribution of subtle motor dysfunction cannot be excluded. Together, these findings support USV emission as a potentially sensitive functional readout of early ALS-related changes.

## Conclusions

Collectively, our results demonstrate that native FL TDP-43 is not a passive component of pathological inclusions but an active driver of toxicity and disease propagation *in vivo*. By showing that a single exposure to FL TDP-43 is sufficient to induce corticospinal spreading, mitochondrial dysfunction, neurodegeneration, and peripheral impairment, this study fills a critical gap in the field. Notwithstanding the invaluable insights provided by human neuropathological studies, such approaches are inherently limited to end-stage material and cannot resolve the temporal sequence of pathological spreading, particularly given the rapid progression of ALS.^64,69^ In this context, the model described here provides a non-transgenic, TDP-43-driven platform that reproduces key pathological and functional features of ALS, including early mitochondrial alterations, region-specific vulnerability, and brain-to-muscle propagation. This system offers a valuable tool to investigate disease mechanisms and to test therapeutic strategies targeting TDP-43 toxicity and spreading.

## Acknowledgements

The authors gratefully acknowledge the financial support from the Italian Ministry of University (PRIN 2020-2020PBS5MJ). The authors also gratefully acknowledge Prof. Angelo Poletti for his valuable contribution to the interpretation of the results.

We acknowledge Cesast (Academic Services Center for Animal Facility) for animal care and housing and Cesar (Academic Services Center for Research Support) for technical assistance at the University of Cagliari. We also acknowledge Dr. Gessica Piras for her skilled assistance throughout the in vivo experimentation.

## Notes

### Competing Interest Statement

The authors have declared no competing interest.

