## Supplemental figures S1-S6 and Table 1 for "Corticospinal propagation of full-length TDP-43 toxicity drives brain-to-muscle pathology"

**Fig. S1**

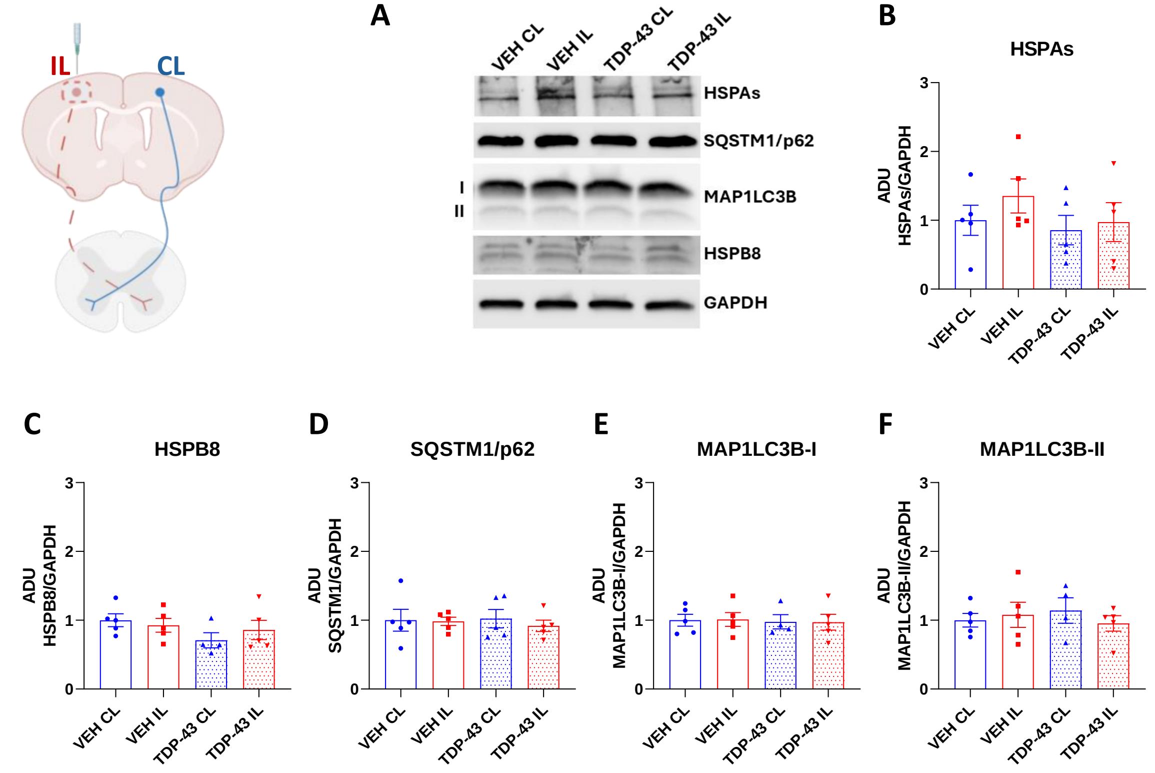

**Fig. S1. TDP-43 infusion does not affect the protein quality control system in the motor cortex.**
**A)** Representative western blot analysis of the chaperone proteins HSPAs (both inducible and constitutive), the small heat shock protein HSPB8, the autophagy receptor SQSTM1/p62 and the autophagy marker MAP1LC3B-I and its lipidated MAP1LC3-II form. **B-F**) Western blot quantifications of HSPAs (**B**), HSPB8 (**C**), SQSTM1/p62, MAP1LC3B-I (**E**) and MAP1LC3B-II (**F**) protein levels. The graphs represent the densitometric analyses of each protein normalized using GAPDH as loading control. Data are expressed as arbitrary densitometric units (ADU). Mean ± SEM of five independent replicates (ANOVA with Tukey's post hoc test among groups). Please note that dotted red columns in the graphs always refer to the TDP-43-affected side (i.e., ipsilateral to the injection side for the motor cortex and contralateral to the injection side for the spinal cord and muscle). CL = contralateral to the injection site; IL = ipsilateral to the injection site; VEH = vehicle.

**Fig. S2**

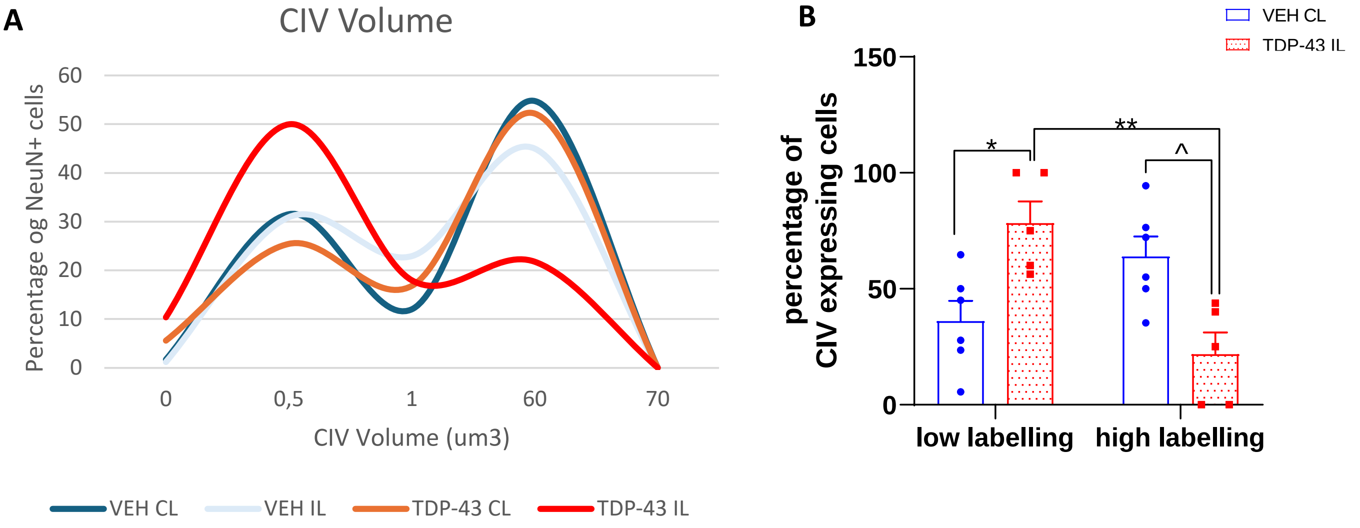

**Fig. S2. A)** Frequency distribution showing CIV colocalization within NeuN+ cells in the motor cortex four-months post infusion. **B)** NeuN+ cells were categorized into two sub-populations, based on the CIV expression, namely low labelled and high labelled. Values represent the mean ± SEM (One-way ANOVA followed by Tukey's post hoc test) *$p<0.05$ vs VEH CL 0-1; ^$p<0.05$ vs VEH CL 2-60; **$p<0.001$ vs TDP-43 IL 0-1. CL=contralateral to the injection site; IL=ipsilateral to the injection site; VEH=vehicle. Please note that dotted red columns in the graphs always refer to the TDP-43-affected side (i.e., ipsilateral to the injection side for the motor cortex and contralateral to the injection side for the spinal cord and muscle). C = complex; CL = contralateral to the injection site; IL = ipsilateral to the injection site; VEH = vehicle.

Fig. S3

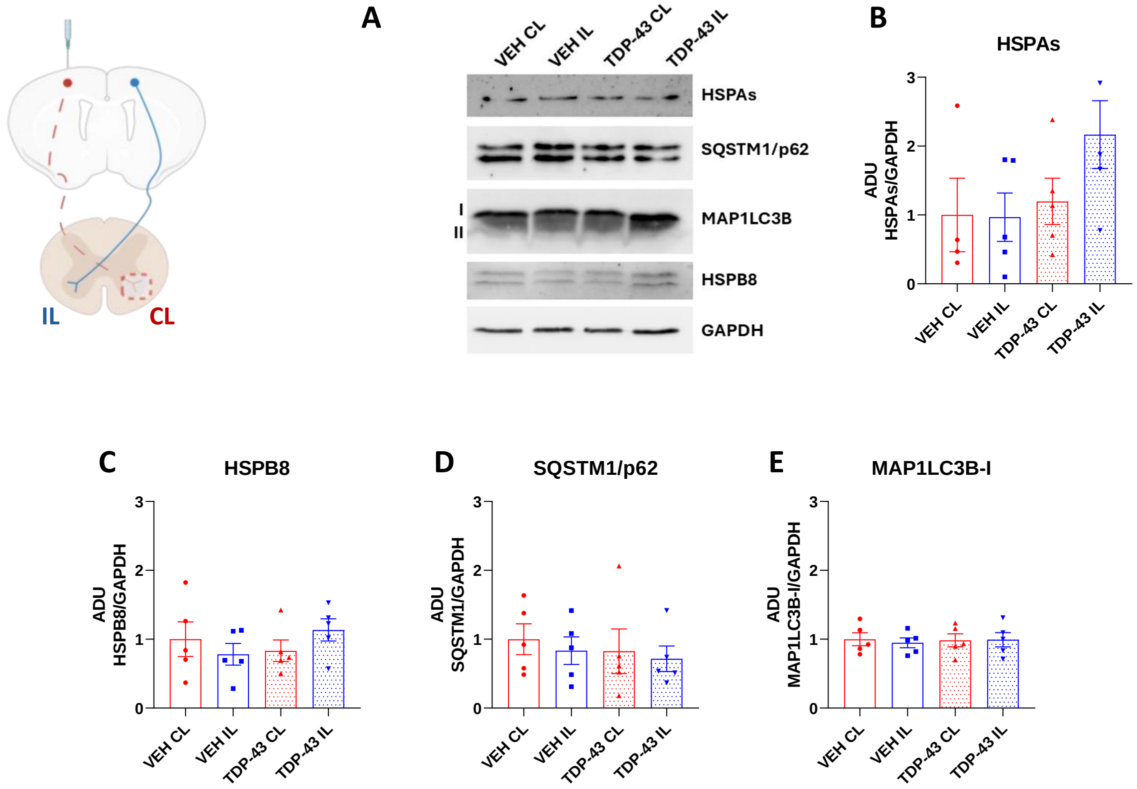

**Fig. S3. TDP-43 infusion doesn't affect the protein quality control system in the cervical tract of the spinal cord.**

**A)** Representative western blot analysis of the chaperone proteins HSPAs (both inducible and constitutive), the small heat shock protein HSPB8, the autophagy receptor SQSTM1/p62 and the autophagy marker MAP1LC3B-I and its lipidated MAP1LC3-II form. **B-E**) Western blot quantifications of HSPAs (**B**), HSPB8 (**C**), SQSTM1/p62 and MAP1LC3B (**E**) protein levels. The graphs represent the densitometric analyses of each protein normalized using GAPDH as loading control. Data are expressed as arbitrary densitometric units (ADU). Mean ± SEM of five independent replicates (ANOVA with Tukey's post hoc test among groups). Please note that dotted red columns in the graphs always refer to the TDP-43-affected side (i.e., ipsilateral to the injection side for the motor cortex and contralateral to the injection side for the spinal cord and muscle). CL = contralateral to the injection site; IL = ipsilateral to the injection site; VEH = vehicle.

Fig. S4

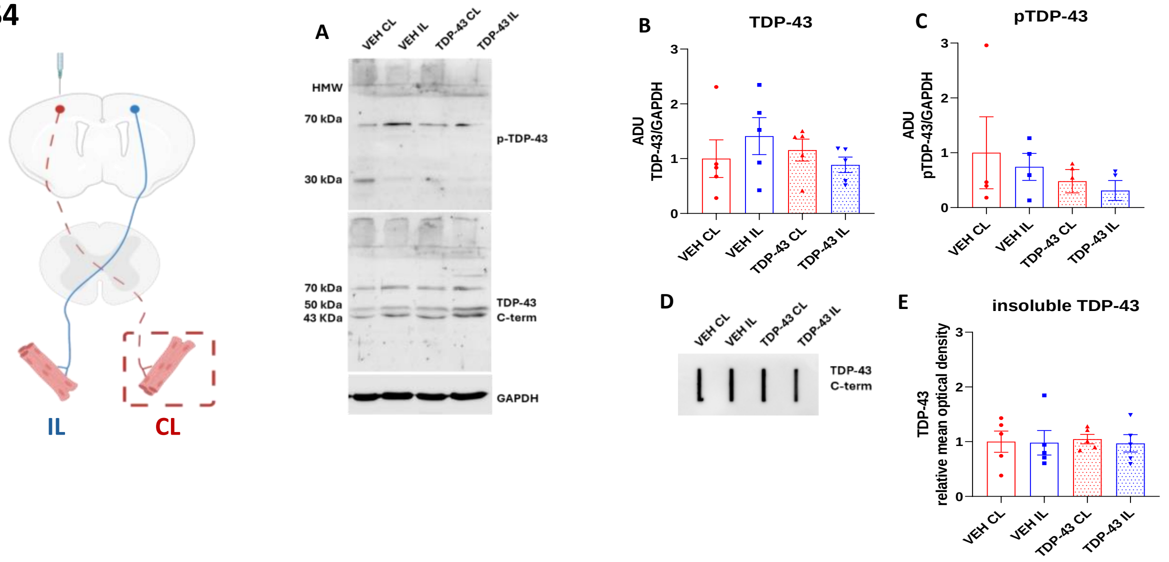

**Fig. S4. TDP-43 behavior in muscle.**

**A-C**) Representative western blot analysis (**A**) and relative quantifications of the C-terminal TDP-43 (**B**) and pTDP-43 (**C**) species. The graphs represent the densitometric analyses of C-terminal and phospho-TDP-43 signals normalized using GAPDH as loading control. Data are expressed as arbitrary densitometric units (ADU). Each bar represents the mean ± SEM of five independent replicates (ANOVA with Tukey's post hoc test among groups). **D-E**) Representative filter retardation assay (**D**) and relative densitometric analysis (**E**) of C-terminal insoluble TDP-43 species. Value represents the mean ± SEM of five independent replicates (ANOVA with Tukey's post hoc test among groups). Please note that dotted red columns in the graphs always refer to the TDP-43 affected side (i.e., ipsilateral to the injection side for the motor cortex and contralateral to the injection side for the spinal cord and muscle). Please note that dotted red columns in the graphs always refer to the TDP-43-affected side (i.e., ipsilateral to the injection side for the motor cortex and contralateral to the injection side for the spinal cord and muscle). CL = contralateral to the injection site; IL = ipsilateral to the injection site; VEH = vehicle.

Fig. S5

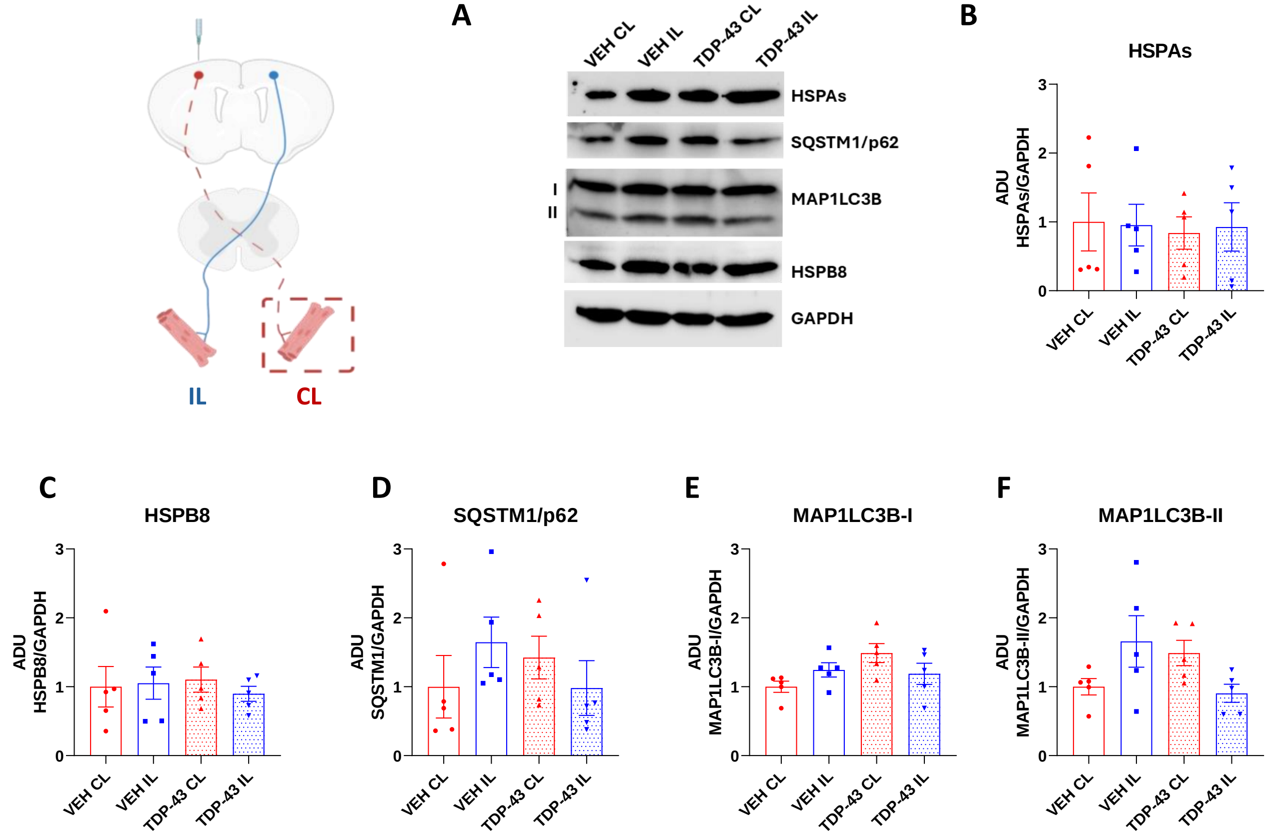

**Fig. S5. TDP-43 infusion doesn't affect the PQC system in the gastrocnemius.**
**A)** Representative western blot analysis of the chaperone proteins HSPAs (both inducible and constitutive), the small heat shock protein HSPB8, the autophagy receptor SQSTM1/p62 and the autophagy marker MAP1LC3B-I and its lipidated MAP1LC3-II form. **B-E**) Western blot quantifications of HSPAs (**B**), HSPB8 (**C**), SQSTM1/p62, and MAP1LC3B-I (**E**) protein levels. The graphs represent the densitometric analyses of each protein normalized using GAPDH as loading control. Data are expressed as arbitrary densitometric units (ADU). Mean ± SEM of five independent replicates (ANOVA with Tukey's post hoc test among groups). Please note that dotted red columns in the graphs always refer to the TDP-43-affected side (i.e., ipsilateral to the injection side for the motor cortex and contralateral to the injection side for the spinal cord and muscle). CL = contralateral to the injection site; IL = ipsilateral to the injection site; VEH = vehicle.

**Fig. S6**

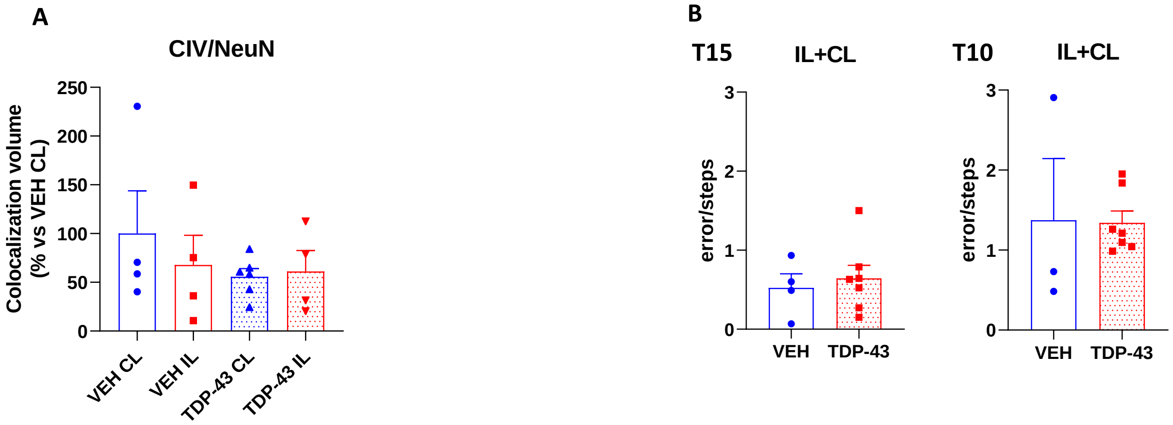

**Fig. S6. Lack of subcellular and behavioral effects of TDP-43 one month post-infusion**. **A)** Total volume occupied by CIV colocalized with $NeuN^+$ cells. **B)** Sensorimotor deficits were assessed using the challenging beam walk test. T15 and T10 correspond to the 15-mm and 10-mm beams, respectively. Graphs represent ipsilateral and contralateral paws relative to the injection site. Values represent mean ± SEM. C = complex; CL = contralateral to the injection site; IL = ipsilateral to the injection site; VEH = vehicle.

**Table 1**.
Uncoupled respiration (DNP 0.2 mM) on gastrocnemius isolated mitochondria, nmolO2/mg/min (data are mean ± SEM).

| Sample | CI | CII | CIV |
|---|---|---|---|
| CL vehicle | 487.00±48.25 | 460.18±31.9* | 1097.00±128.02 |
| IL vehicle | 522.37±48.88 | 488.96±40.06 | 958.00±125.76 |
| CL TDP-43 | 469.92± 62.54 | 557.58±38.47* | 989.33±145.54 |
| IL TDP-43 | 565.89±67.55 | 579.31±51.52 | 1017.02±65.28 |

*p= 0.06 Mann–Whitney test TDP-43 versus vehicle

**Table 2.**
State 3, state 4 (both nmolO2/mg/min), RCR, and ADP/O in isolated mitochondria from vehicle and TDP-43 treated CL gastrocnemius (data are mean ± SEM).

| Sample | CI | CII | Lipid substrate |
|---|---|---|---|
| CL vehicle | | | |
| State 3 | 232.03±30.9 | 412.31±34.31 | 124.85±11.19 |
| State 4 | 31.72±3.45 | 128.13±11.32 | 30.53±2.84 |
| RCR | 7.7±0.75 | 3.25±0.12 | 4.27±0.48 |
| ADP/O | 2.85±0.23 | 1.16±0.28 | 2.28±0.17 |
| CL TDP-43 | | | |
| State 3 | 277.64±32.7 | 423.87±27.28 | 155.73±0.12 |
| State 4 | 37.15±3.82 | 137.8±9.07 | 33.88±2.85 |
| RCR | 7.56±0.6 | 3.12±0.17 | 4.79±0.24 |
| ADP/O | 2.62±0.24 | 0.92±0.15 | 2.22±0.24 |
